# Proinflammatory Cytokines Suppress Nonsense-Mediated RNA Decay to Impair Regulated Transcript Isoform Processing in Pancreatic β-Cells

**DOI:** 10.1101/2023.12.20.572623

**Authors:** Seyed. M. Ghiasi, Piero Marchetti, Lorenzo Piemonti, Jens H. Nielsen, Bo T. Porse, Thomas Mandrup-Poulsen, Guy A. Rutter

**Author notes:** Lead investigator (Seyed. M. Ghiasi). equally contributing to this study (Thomas Mandrup-Poulsen, Guy A. Rutter).

## Abstract

Proinflammatory cytokines are implicated in pancreatic β-cell failure in type 1 and type 2 diabetes and are known to stimulate alternative RNA splicing and the expression of Nonsense-Mediated RNA Decay (NMD) components. Here, we investigate whether cytokines regulate NMD activity and identify transcript isoforms targeted in β-cells. A luciferase-based NMD reporter transiently expressed in rat INS1(832/13), human-derived EndoC-βH3 or dispersed human islet cells is used to examine the effect of proinflammatory cytokines (Cyt) on NMD activity. Gain- or loss-of function of two key NMD components UPF3B and UPF2 is used to reveal the effect of cytokines on cell viability and function. RNA-sequencing and siRNA-mediated silencing are deployed using standard techniques. Cyt attenuate NMD activity in insulin-producing cell lines and primary human β-cells. These effects are found to involve ER stress and are associated with downregulation of UPF3B. Increases or decreases in NMD activity achieved by UPF3B overexpression (OE) or UPF2 silencing, raises or lowers Cyt-induced cell death, respectively, in EndoC-βH3 cells, and are associated with decreased or increased insulin content, respectively. No effects of these manipulations are observed on glucose-stimulated insulin secretion. Transcriptomic analysis reveals that Cyt increase alternative splicing (AS)-induced exon skipping in the transcript isoforms, and this is potentiated by UPF2 silencing. Gene enrichment analysis identifies transcripts regulated by UPF2 silencing whose proteins are localized and/or functional in extracellular matrix (ECM) including the serine protease inhibitor SERPINA1/α-1-antitrypsin, whose silencing sensitises β-cells to Cyt cytotoxicity. Cytokines suppress NMD activity via UPR signalling, potentially serving as a protective response against Cyt-induced NMD component expression. Our findings highlight the central importance of RNA turnover in β-cell responses to inflammatory stress.

## Introduction

Inflammatory and glucolipotoxic (GLT) stress causing β-cell failure and destruction *in vitro* differentially regulates hundreds of β-cell transcripts (1, 2). The upregulation of splicing factors and of proteins involved in pre-mRNA processing gives rise to alternative splicing (AS) events, which in turn deregulates the balance and turnover of transcript isoforms (3). Interestingly, most human mRNAs exhibit alternative splicing, but not all alternatively spliced transcripts are translated into functional proteins and are therefore targeted for degradation via the RNA decay pathways.

In addition to regulating the expression of normal transcripts, the human nonsense-mediated RNA decay (NMD) machinery functions to eliminate premature termination codon (PTC)-containing mRNAs, as reviewed extensively (4). Alternatively, spliced mRNA species and translation of dominant transcript isoforms vary in a cell-specific manner and depends on the capacity of cells to cope with damaged transcripts (5–7). A substantial number (i.e., around 35%, depending on tissue and physiological conditions) of alternatively spliced variants contain a PTC (4, 8, 9). Approximately 35% of the cytokines-regulated transcripts in human islets undergo alternative splicing (6), and Cyt profoundly up-regulate NMD) in rat and human insulin-producing cell lines and primary β-cells, likely to handle the NMD-load inferred by PTC containing splice variants (4, 10, 11).

However, in addition to canonical NMD in which all key NMD components function on target transcripts, a second branch of NMD is (in)dependently regulated in an autoregulatory feedback loop by its key factors including UPF2 and UPF3 in a cell type-specific manner as reviewed previously (4, 10).

In a previous study (11) we profiled the *expressional level* of NMD components and their regulation by cytokines and GLT in insulin-producing cells, but the NMD *activity* and its consequences for the β-cell transcriptome remained to be investigated. Here, using a luciferase-based NMD activity reporter, gain-/loss-of function and RNA-sequencing analyses in rodent and human β-cell systems, we measured NMD activity and explored its consequences for function and viability of pancreatic β-cells under normal conditions and inflammatory stress.

## Materials and Methods

### Cell culture, human islet dispersion and treatment

INS1(832/13) (12), EndoC-βH3 (13) or dispersed human islet cells were cultured and manipulated according to the protocols and procedures described in Supplementary Methods.

### Luciferase-based NMD activity assay

One million cells were co-transfected with 650 ng of plasmid encoding either human *Haemoglobin-β* (*HBB*) wildtype (WT or PTC-) or with a PTC-containing mutation (NS39 or PTC+) fused with *Renilla (RLuc),* in brief named PTC-and PTC+), respectively. *Firefly (FLuc)* plasmid (14) was used as transfection efficiency reference. *Renilla* and *Firefly* luminescence was measured by Dual-Luciferase Reporter Assay (Promega, Hampshire, England) (Supplementary Methods). *RLuc* signals were normalized to the *FLuc* control in both HBB NS39, in the following named HBB(PTC+) and HBBWT, in the following named HBB (PTC-), and NMD activity was calculated by dividing the *RLuc/FLuc*-HBB PTC-) by the *RLuc/FLuc-*HBB PTC+) (Supplementary Fig.1A) (14). Experiments where the control construct *RLuc/FLuc*-HBB(PTC-) was affected by cytokines were excluded, so that the resulting NMD activity only denotes the PTC-containing HBB( PTC+). The transfection efficiency was tested twice and resulted in an average of 80% in INS1 and EndoC-βH3 cells as measured by FACS analysis of cells transfected with a GFP expressing plasmid (Supplementary Fig.1B-C).

### Functional analysis of UPF3A/B overexpression

One million INS1(832/13) or EndoC-βH3 cells were transfected with 650 ng of plasmids encoding UPF3A, UPF3B or UPF3BΔ42 (15), then simultaneously with NMD activity reporter plasmids (as above, 650 ng/million cells), recounted and seeded for Western blotting, glucose-stimulated insulin secretion (GSIS), viability, apoptosis (detailed below) and NMD activity assays in relevant plates and pre-incubated for 48 h before treatment with cytokines as explained in the Supplementary Methods.

### Lentiviral shRNA gene knock-down

GPIZ lentiviral shRNAs particles directed against *UPF2, Upf3A* or *Upf3B*, and a non-silencing shRNA (NS) as negative control, were produced using the Trans-Lentiviral shRNA Packaging System in HEK293 cells (Horizon, Cambridge, England) according to the manufacturer’s protocol (Supplementary Methods).

### Apoptosis and cell viability assays

Apoptosis assays were performed in duplicate by detection of caspase-3 activity using a fluorometric [µM AMC] (or/colorimetric [µM PNA/min/ml] unless stated) assay kit (Cat#APPA015-1KT/CASP3C-1KT, Sigma, London, England) according to the manufacturers’ protocols. Cell viability was measured by Alamarblue assay (Cat#DAL1025, LifeTechnologies, Renfrew, England) as previously described (11).

### Library preparation, RNA-sequencing, and data analysis

Thirty-three independent biological replicates of total RNA from the NS control and or UPF2 KD EndoC-βH3 cells exposed to cytokines, GLT, or PBS (i.e., N=6 of each PBS-/or cytokines-exposed NS control and UPF2 KD, and N=4/N=5 of GLT-exposed NS control/UPF2 KD, respectively) was extracted using TRIZOL, treated with DNase, and precipitated with isopropanol (Supplementary Methods). One µg total RNA/per isolate was used as input for generation of sequencing libraries using NEBNext®Ultra-TM RNA Library-Prep (Cat#E7770, NEB, Ipswich, USA) following manufacturer’s recommendations (Supplementary Methods). The RNA-seq raw data underwent quality control were mapped to human reference genome (16) and analysed using the bioinformatic pipeline described in the Supplementary Methods.

### cDNA synthesis and RT-qPCR

Purified total RNA (500 ng) was used for cDNA synthesis with the SuperScript™ (Cat# 11904018, LifeTechnologies). Real-time Reverse Transcriptase-quantitative PCR (RT-qPCR) was performed on 12 ng cDNA with SybrGreen PCR mastermix (LifeTechnologies) and specific primers (supplementary Table 2) and run in an ABI? real-time PCR machine (Applied Biosystems, ThermoFisher Scientific). The raw data was analysed through -ΔCt as described in Supplementary Methods.

### Western blotting

Western blotting was performed using antibodies against alpha-Tubulin (1:2000) (Cat# T5168, Sigma), UPF2 (1:1000) (Cat# PA5-77128, LifeTechnologies), Upf3A (1:1000) (Cat# PA5-41904, LifeTechnologies), UPF3B (1:1000) (Cat#PB9843, Boster-Bio, Pleasanton, USA) and α-1-antitrypsin (1:1000) (Cat#TA500375, LifeTechnologies) as described in Supplementary Methods (17).

### Glucose-stimulated insulin secretion (GSIS)

Three hundred thousand INS1(832/13) or EndoC-βH3 cells were cultured in 12-well plates (Cat#150200, Nunc, Buckingham, England), and pre-incubated for two days. GSIS was carried out using Krebs-Ringer buffer containing 2 mM or 17 mM glucose as described (11, 18).

### Insulin assay

Insulin concentration (ng/ml or pM) was measured using rat insulin ultra-sensitive ELISA kit (Cat#62IN2PEG, Cisbio, Cambridge, England) or human insulin ELISA kit (Cat#90095, CrystalChem, IL, USA), respectively, according to manufacturer’s protocol.

### Statistical analysis

Data are presented as means ± SEM. Statistical analysis was carried out on raw data also in cases where figures give normalized data. Group comparisons were carried out by two- or one-way ANOVA as appropriate. Significant ANOVAs were followed by post-hoc paired Student’s t-test with Bonferroni-correction using GraphPad Prism 6.0 (La Jolla, USA). Paired t-test was chosen to normalize for inter-passage variability in outcome parameters. Since the experimental conditions did not allow sequential sampling from the same cell culture, parallel control and interventional plate wells were considered to be paired observations and analysed accordingly statistically. If the *post-hoc* paired t-test did not reveal a carrying statistical difference by ANOVA, individual paired t-tests were performed and corrected for multiple comparisons. Bonferroni-corrected *P*-values ≤0.05 were considered significant and ≤0.10 a trend.

## Results

### Cytokines suppress NMD activity in β-cells

We previously reported that cytokines and glucolipotoxicity differentially up or down-regulate NMD component transcripts in pancreatic β-cells (11). However, whether this regulation leads to increased NMD *activity* remained to be elucidated. Here, we used a luciferase-based NMD reporter (Supplementary Fig.1A) (14) to examine NMD activity in rat INS1(832/13), human insulin-producing EndoC-βH3 cells and primary human islets. Luciferase activity analysis showed that cytokines (Cyt;150 pg/mL IL-1β +0.1 ng/mL IFNγ+0.1 ng/mL TNFα) significantly suppressed NMD activity by nearly 50% after 18 h, but not 6h, of exposure in INS1(832/13) cells (Supplementary Fig.2A-i and iii).

We then tested the effects of cytokines on EndoC-βH3 cells and dispersed human islet cells. Cytokines (2.5 ng/ml IL-1β+10 ng/ml TNFα+10 ng/ml IFNγ, chosen from dose-response experiment shown in Supplementary Fig.2B-i) attenuated NMD activity by 30% (*p=*0.009, n=6) and 40% (*p=*0.0006, n=6) after 18 h exposure of EndoC-βH3 cells (Fig.1A-i) and dispersed human islet cells (Fig.1B), respectively. Cyt increased the luciferase signal (*RLuc/FLuc*) from the HBB(PTC+) (Supplementary Fig.2B-ii, Supplementary Fig.2C) but not from the HBB(PTC+) (Supplementary Fig. 2B-v and 2C-ii) in both cell models confirming that the NMD substrate HBB(PTC+) was restored due to NMD activity attenuation by Cyt.

**Figure 1.**
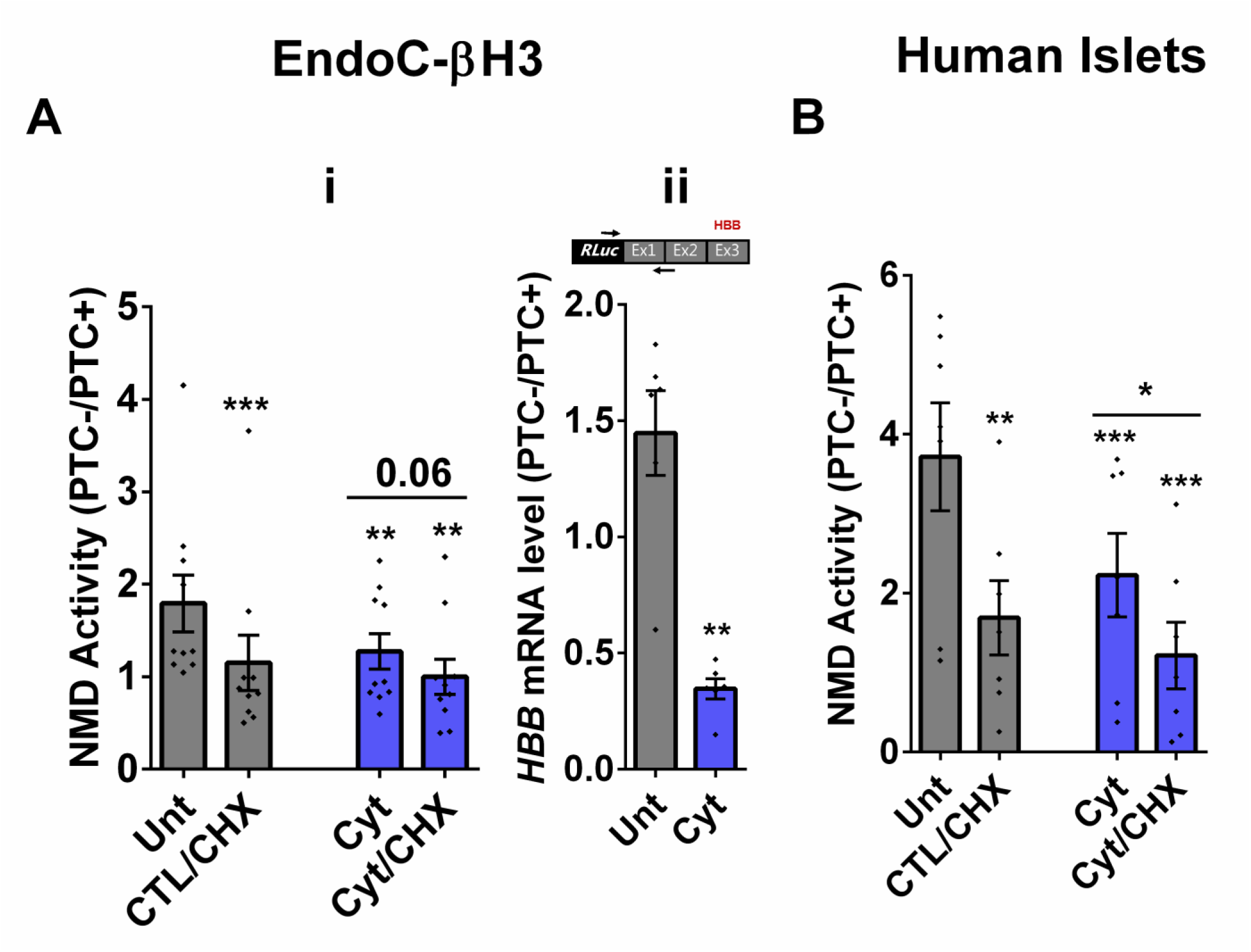
Cytokines suppress NMD activity in β-cells. **A-B**. EndoC-βH3 cells (A) and dispersed human islet cells (B) were co-transfected with *Renilla*-HBB(WTnamed PTC-) and or *Renilla*-HBB(NS39 (named PTC+), and the *Firefly* plasmids and exposed to cytokines combination and or PBS as untreated (Unt) simultaneously with or without Cycloheximide (CHX) as positive control for inhibited NMD activity. Luciferase activity was measured in the lysate of the transfected EndoC-βH3 cells (A-i) and dispersed human islet cells (B) exposed to cytokines combination (Cyt; 3 ng/mL IL-1β + 10 ng/mL IFNγ+ 10 ng/mL TNFα) for 18 hrs. A-ii. mRNA level of *Renilla-HBB* fused gene and *Firefly* gene in the transfected EndoC-βH3 cells was quantified by RT-qPCR using specific primers extending the junction of exons 1 and 2 of the *HBB* gene, and *Renilla* gene, or only *Firefly* gene and normalised to actin and tubulin, respectively. The data are means ± SEM of N=6. The symbol * indicates the Bonferroni-corrected paired t-test values of treatments versus untreated (Unt) (A-B) or otherwise, cytokines (Cyt) that is designated by a line on top of the bars (A-B). * ≤ 0.05, ** ≤ 0.01, *** ≤0.001, **** ≤ 0.0001. ns: non-significant. HBB: *Haemoglobin-β,* PTC: premature termination codon. On the top of Fig. 1A-i, RLuc: *Renilla* Luciferase and Ex: exon.

We next examined whether cytokines-mediated suppression of NMD was consistent with an accumulation of HBB(PTC+) *transcripts*. For this, we used a forward and reverse primer set to amplify the *Renilla* gene and the junction of exons 1 and 2 (i.e., ensuring amplification of mature transcripts only), respectively. RT-qPCR analysis demonstrated that cytokines caused significant up-regulation of HBB(PTC+), but not HBB(PTC-) mRNA levels, rendering a significant reduction of the relative PTC-/PTC+ mRNA levels in INS1(832/13) (*p=*0.008) (Supplementary Fig.2A-v and vi) and EndoC-βH3 (*p=*0.001) cells (Fig.1A-ii; Supplementary Fig.2B-iv), which verified the suppressive effect of cytokines on NMD activity.

We also examined the effect of 25 mM glucose or GLT conditions on the NMD activity. Unfortunately, these conditions significantly affected the luciferase signal (*RLuc/FLuc*) from the HBB(PTC-) control, preventing further study of the effects of metabolic stressors on NMD activity.

Taken together, these results show that cytokines suppress the activity of the NMD activity in a range of insulin secreting cell types.

### Cytokines-induced suppression of NMD activity in β-cells is ER stress dependent

Whereas NMD degrades unfolded protein response (UPR)-induced transcripts in compensated ER stress, NMD is suppressed in response to pronounced endoplasmic reticulum (ER) stress to allow a full-blown UPR (19, 20). Cytokines induce a robust ER stress in pancreatic β-cells, largely via nuclear factor-κB (NF-κB) activation and production of nitroxidative species that inhibit the smooth endoplasmic reticulum Ca^2+^ ATPase (SERCA) 2B pump, leading to ER calcium depletion (11, 17). We have previously shown that chemical inhibition of inducible nitric oxide synthase (iNOS) alleviated ER stress and normalised cytokines-mediated regulation of NMD components in INS1 cells (11). Therefore, we asked, if cytokines-mediated reduction of NMD activity was dependent on an ER stress response in β-cells. We first demonstrated that thapsigargin (TG), a non-competitive inhibitor of SERCA (21) and ER stress inducer (22) inhibited NMD activity by 50% in EndoC-βH3 cells as measured by luciferase assay (Supplementary Fig.3A-i and ii). Compared to untreated EndoC-βH3 cells (Unt), cytokines significantly augmented the increase in mRNA levels encoding the ER stress markers BiP, Xbp1 and Chop (FDR <0.05) measured by RNA-sequencing analysis (Fig.2A-i), and later verified by RT-qPCR examination (Fig.2A-ii-vi). Finally, compared with Unt, cytokines significantly decreased the NMD activity by 30%, and this effect was counteracted by the protein kinase R-like endoplasmic reticulum kinase (PERK) phosphorylation inhibitor GSK157 (8 µM) and by the Inositol-Requiring Enzyme1 (IRE1α) endoribonuclease inhibitor 4µ8C (16 µM) in EndoC-βH3 cells (Fig.2B-i, Supplementary Fig.3B and Supplementary Fig.3C-i and ii). RT-qPCR analysis of the relative PTC-/PTC+ mRNA levels in EndoC-βH3 cells confirmed the NMD activity data (Fig.2B-ii, Supplementary Fig.3C-iii).

**Figure 2.**
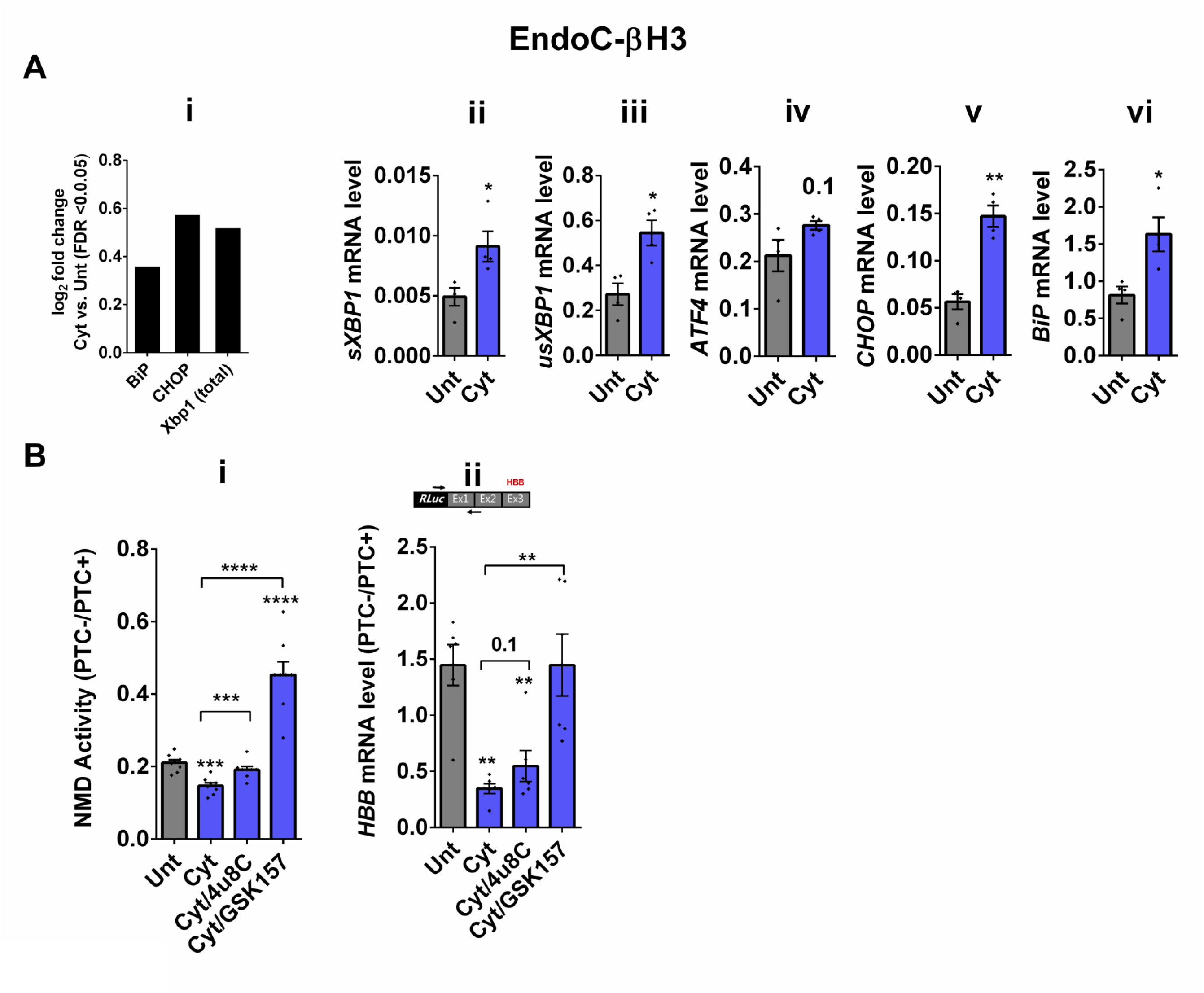
Cytokines-induced suppression of NMD activity in β-cells is ER stress dependent. A. mRNA levels of ER stress markers in EndoC-βH3 cells exposed to cytokines combination (Cyt; 3 ng/mL IL-1β + 10 ng/mL IFNγ+ 10 ng/mL TNFα) for 18 hrs was quantified by RNA-sequencing (A-i) with false discovery rate (FDR) <0.05 presented as logarithmic fold change the cytokines (Cyt) treatment versus control (untreated), and RT-qPCR (A-ii, iii, iv and v) which was normalised to tubulin mRNA. B. EndoC-βH3 cells were co-transfected with *Renilla*-HBB (PTC-) and or *Renilla*-HBB (PTC+), and the *Firefly* plasmid and exposed to PBS as untreated (Unt), cytokines combination (Cyt; 3 ng/mL IL-1β + 10 ng/mL IFNγ+ 10 ng/mL TNFα) alone, and or simultaneously with 16 µM of 4μ8C, an endoribonuclease inhibitor of IRE1α, and or 8 µM of GSK2656157 (GSK157), PERK inhibitor for 18 hrs. B-i. Luciferase activity was measured in the lysate of EndoC-βH3 cells transfected with *Renilla*-HBB(PTC-) and or *Renilla*-HBB(PTC+), and the *Firefly* plasmid exposed to PBS as untreated (Unt) or given conditions, and represented as NMD activity calculated by dividing luciferase activity of HBB(PTC-)/HBB(PTC+) as explained in the methods. B-ii. mRNA level of *Renilla-HBB* fused gene and *Firefly* gene in the EndoC-βH3 cells was quantified by RT-qPCR using specific primers extending the junction of exons 1 and 2 of the *HBB* gene, and *Renilla* gene, or only *Firefly* gene and normalised to tubulin. The data are means ± SEM of N=6. The symbol * indicates the Bonferroni-corrected paired t-test values of treatments versus untreated (Unt) (A-B) or, otherwise, cytokines (Cyt) that is designated by a line on top of the bars (B). * ≤ 0.05, ** ≤ 0.01, *** ≤0.001, **** ≤ 0.0001. FDR: False discovery rate. On the top of Fig. 2B-right Subpanel, RLuc: *Renilla* Luciferase and Ex: exon.

Taken together, these results demonstrate that inhibition of UPR antagonises the cytokine-mediated reduction of NMD activity in EndoC-βH3, indicating that cytokine-mediated inhibition of NMD activity is UPR-dependent.

### Cytokines-induced suppression of NMD activity is associated with UPF3B downregulation and attenuated by UPF3 overexpression in β-cells

Since we observed in our previous study that cytokines-induced ER stress downregulated UPF3B expression in human and rodent β-cells, as recovering nitroxidative-driven ER stress using the inducible nitric oxide synthase (iNOS) inhibitor N-methyl-l-arginine (NMA) (11), since transcripts encoding UPR components are NMD targets and have been shown to be stabilised by UPF3A/B depletion (19) and since UPF3B is a NMD activator in mammalian cells (23), which led to the proposal of Upf3-dependent and -independent branches of the NMD pathway (4, 10, 24). we reasoned that UPF3 regulated NMD activity in β-cells. We therefore first measured the UPF3A/B expression level and next investigated the functional impact of overexpressing UPF3A/B on cytokines-mediated suppression of NMD activity in β-cells. RT-qPCR examination showed that cytokines significantly downregulated UPF3B mRNA levels after 18 h in both EndoC-βH3 (Fig.3A-i and ii) and INS1(832/13) (Supplementary Fig.4A-i and ii) as previously reported (11). Immunoblot analysis verified overexpression of UPF3A, UPF3B and the UPF3B dominant negative UPF3BΔ42 in both EndoC-βH3 (Supplementary Fig.4B-i) and INS1(832/13) (Supplementary Fig.4C-i). Cytokines reduced NMD activity and overexpression of UPF3B significantly attenuated this reduction in EndoC-βH3 (Fig.3B and Supplementary Fig.4B-ii) and to a lesser extent in INS1(832/13) (Supplementary Fig.4C-ii and iii). Neither UPF3A nor UPF3BΔ42 overexpression counteracted cytokines-attenuated NMD activity.

**Figure 3.**
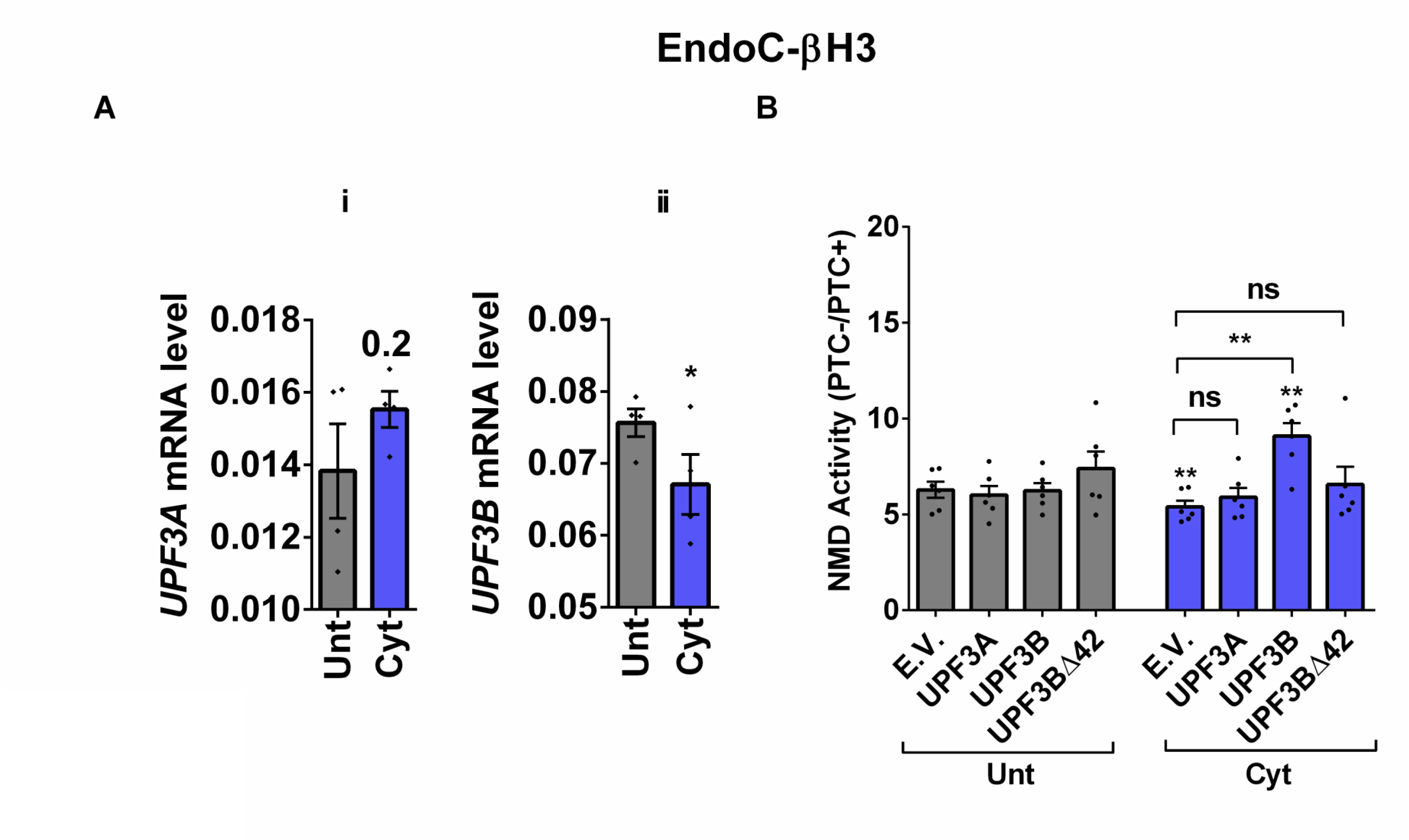
Cytokines-induced suppression of NMD activity is associated with UPF3B downregulation and attenuated by UPF3 overexpression in β-cells. EndoC-βH3 cells were co-transfected with empty vector (E.V.), UPF3A, UPF3B and or UPF3BΔ42 (dominant negative of UPF3B) plasmids, and then with *Renilla*-HBB(PTC-) and or *Renilla*-HBB(PTC+), along with the *Firefly* plasmid and exposed to cytokines combination (Cyt; 3 ng/mL IL-1β + 10 ng/mL IFN-γ+ 10 ng/mL TNFα) for 18 hrs.A-i and ii. mRNA level of *Upf3A* and *Upf3B* genes in EndoC-βH3 cells was quantified by RT-qPCR and normalised to tubulin mRNAs. B. Luciferase activity was measured in the lysate of the transfected cells and represented as NMD activity calculated by dividing luciferase activity of HBB(PTC-)/HBB(PTC+) as explained in the methods. The overexpression of UPF3A and UPF3B proteins was examined by Western blot analysis (Supplementary Fig.4B-i). The data are means ± SEM of N=6. The symbol * indicates the Bonferroni-corrected paired t-test values of treatments versus untreated E.V. (Unt) (A-B) or, otherwise, cytokines (Cyt)-treated E.V. that is designated by a line on top of the bars (B). * ≤ 0.05, ** ≤ 0.01. ns: non-significant.

This result suggests that cytokines reduce the NMD activity in β-cells through downregulation of UPF3B expression.

### UPF3 overexpression deteriorates cell viability and reduces insulin content, but not secretion in EndoC-βH3 cells

The above findings provide evidence that the UPF3-dependent branch of NMD is involved in cytokines-mediated suppression of NMD activity. Therefore, we next investigated the impact of UPF3A/B overexpression on cytokines-induced cell death and insulin secretion. While UPF3A or UPF3B over-expression increased basal cell death, it also exacerbated the cytokines-induced apoptosis in EndoC-βH3 cells (Fig.4A-i and ii). In INS1(832/13) cells neither UPF3A nor UPF3B overexpression changed cell viability in the absence of Cyt exposure, but UPF3B over-expression significantly aggravated cytokines-induced cell death as measured by Alamarblue and caspase-3 activity assays (Supplementary Fig.5A-i and ii). Therefore, we next explored the impact of UPF3A or UPF3B deficiency on β-cell viability. Lentiviral shRNA-mediated knockdown of UPF3A and or UPF3B (Supplementary Fig.5B-i and ii) significantly reduced basal INS1(832/13) cell viability (Supplementary Fig.5C-i-iv). Taken together, this data indicates that genetic manipulations of UPF3A/B could be possibly detrimental for the β-cell viability.

**Figure 4.**
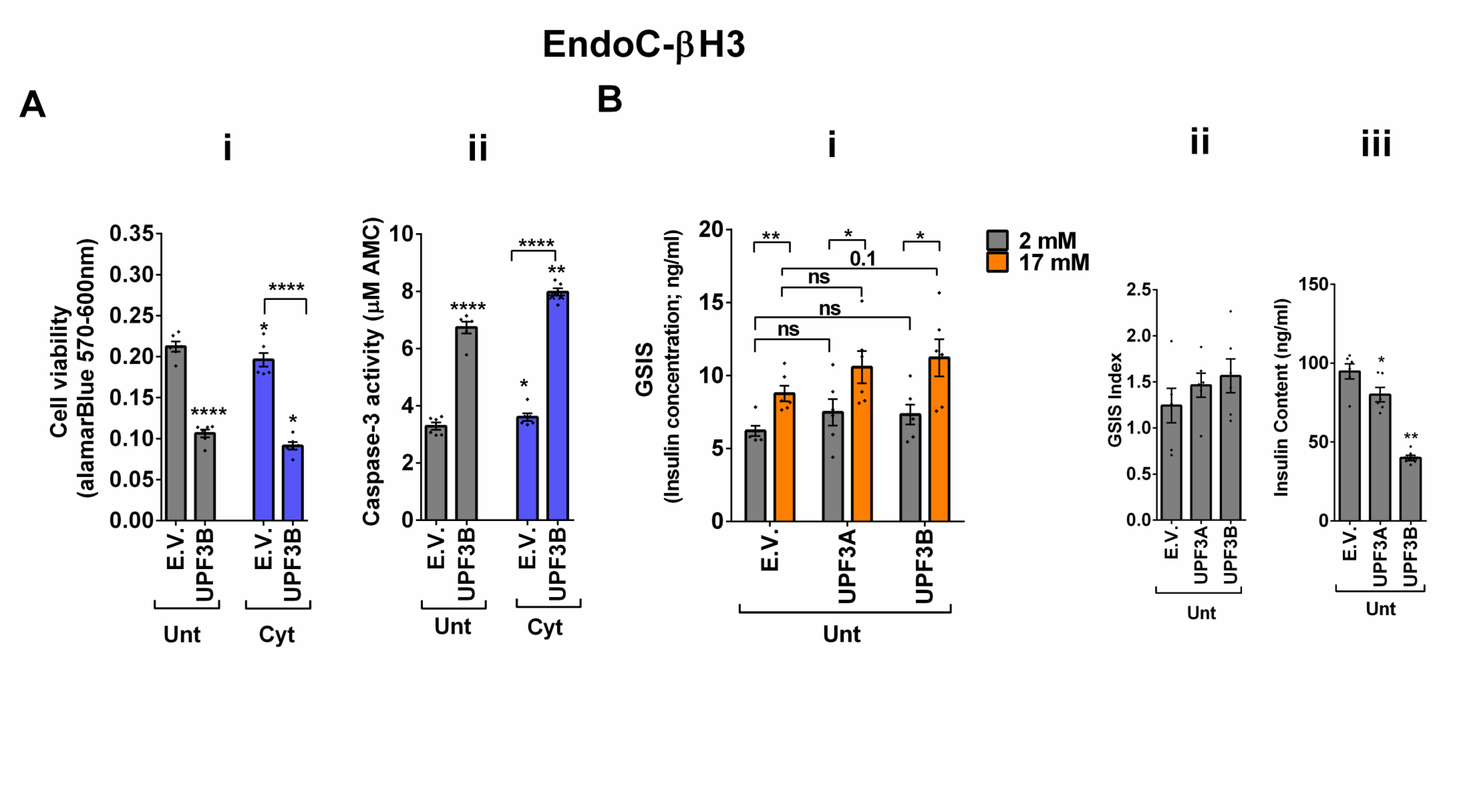
UPF3 overexpression deteriorates cell viability and reduces insulin content, but not secretion in EndoC-βH3 cells. EndoC-βH3 cells were co-transfected with empty vector (E.V.), UPF3A and or UPF3B plasmids and exposed to cytokines combination (Cyt; 3 ng/mL IL-1β + 10 ng/mL IFNγ+ 10 ng/mL TNFα) for three days. A. Cell viability was measured by Alamarblue (A-i) and caspase-3 activity (A-ii) assays (N=6). B. Glucose-stimulated insulin secretion (GSIS) (B-i) and insulin contents (B-iii) were investigated in the transfected EndoC-βH3 cells. Insulin concentration (ng/ml) was measured by insulin ultra-sensitive assay (N=6). GSIS index (B-ii) was calculated by dividing insulin concentration measured in the treatments of 17 mM by 2 mM glucose. The data are means ± SEM of N=6. The symbol * indicates the Bonferroni-corrected paired t-test values of treatments versus untreated E.V. (Unt) or, otherwise, cytokines (Cyt)-treated E.V. that is designated by a line on top of the bars (A), or the Bonferroni-corrected paired t-test values of corresponding low versus high glucose, that is otherwise, designated by lines on the top of the bars (B-i). * ≤ 0.05, ** ≤ 0.01, **** ≤ 0.0001. ns: non-significant.

In contrast to Cyt that downregulate UPF3B in INS1 (11), EndoC-βH3 (Fig.3A-ii) and INS1(832/13) cells (Supplementary Fig.4A-ii), GLT does not downregulate UPF3B (11).We therefore explored the effect of UPF3 deficiency on glucolipotoxicity-induced cell death in β-cells. Measurements of Caspase-3 activity demonstrated that both UPF3A and UPF3B knockdown rendered a slight, but significant protection against 24 h glucolipotoxicity in INS1(832/13) cells in comparison with untreated cells (Supplementary Fig.5D-i and ii). Further, treatment with the NMD activator Tranilast dose-dependently sensitised to glucolipotoxicity-, but not cytokines-induced, EndoC-βH3 cell death, measured by caspase-3 activity assay (Supplementary Fig.4E-i and ii). Neither UPF3A nor UPF3B overexpression affected GSIS in EndoC-βH3 (Fig.4B-I and ii) or INS1(832/13) (Supplementary Fig.5F-I and ii) cells. Nonetheless, UPF3B overexpression profoundly lowered insulin content in EndoC-βH3 cells (Fig.4B-iii). In contrast, knockdown of UPF3A or UPF3B significantly decreased the stimulatory index, as well as provoking a substantial increase in insulin content in control INS1(832/13) cells (Supplementary Fig.5G-ii and iii, and Supplementary Fig.5H-ii and iii), without altering ins1 or 2 mRNA expression (Supplementary Fig.5 G-iv and v, Supplementary Fig.5 H iv and v).

Taken together, these findings reveal that UPF3 overexpression induces basal cell death and exacerbates cytokines-mediated toxicity in β-cells.

### UPF2 knockdown potentiates cytokines suppression of NMD activity and slightly alleviates cytokines toxicity for cell viability and insulin content in EndoC-βH3 cells

Next, we investigated the effect of UPF2 deficiency on the viability and insulin secretion of β-cells because (i) UPF3A and UPF3B are involved in regulating UPF2, a key core NMD activator in mammalian cells (25, 26) by sequestering away from and bridging the exon-junction complex (EJC) with UPF1 and UPF2, respectively, leading to the NMD activation (23), and genome-wide association data (GWAS) data reveal that the *UPF2* variant rs145580445 is significantly associated with type 2 diabetes risk (7). We therefore knocked down the *UPF2* gene in EndoC-βH3 cells using RNA interference and chose the three cell lines in which UPF2 was most efficiently knocked down (KD) (Supplementary Fig.6A-i and ii). Examination of NMD activity using the luciferase-based NMD reporter revealed that UPF2 KD profoundly reduced NMD activity in untreated and cytokines-treated EndoC-βH3 cells (Supplementary Fig.6B-i and ii). Compared with NS control, UPF2 KD slightly, but significantly prevented cytokines-induced cell death (Fig.5A-i and ii). UPF2 KD had no effect on the GSIS, but significantly increased insulin content (Fig.5B-i, ii and iii).

**Figure 5.**
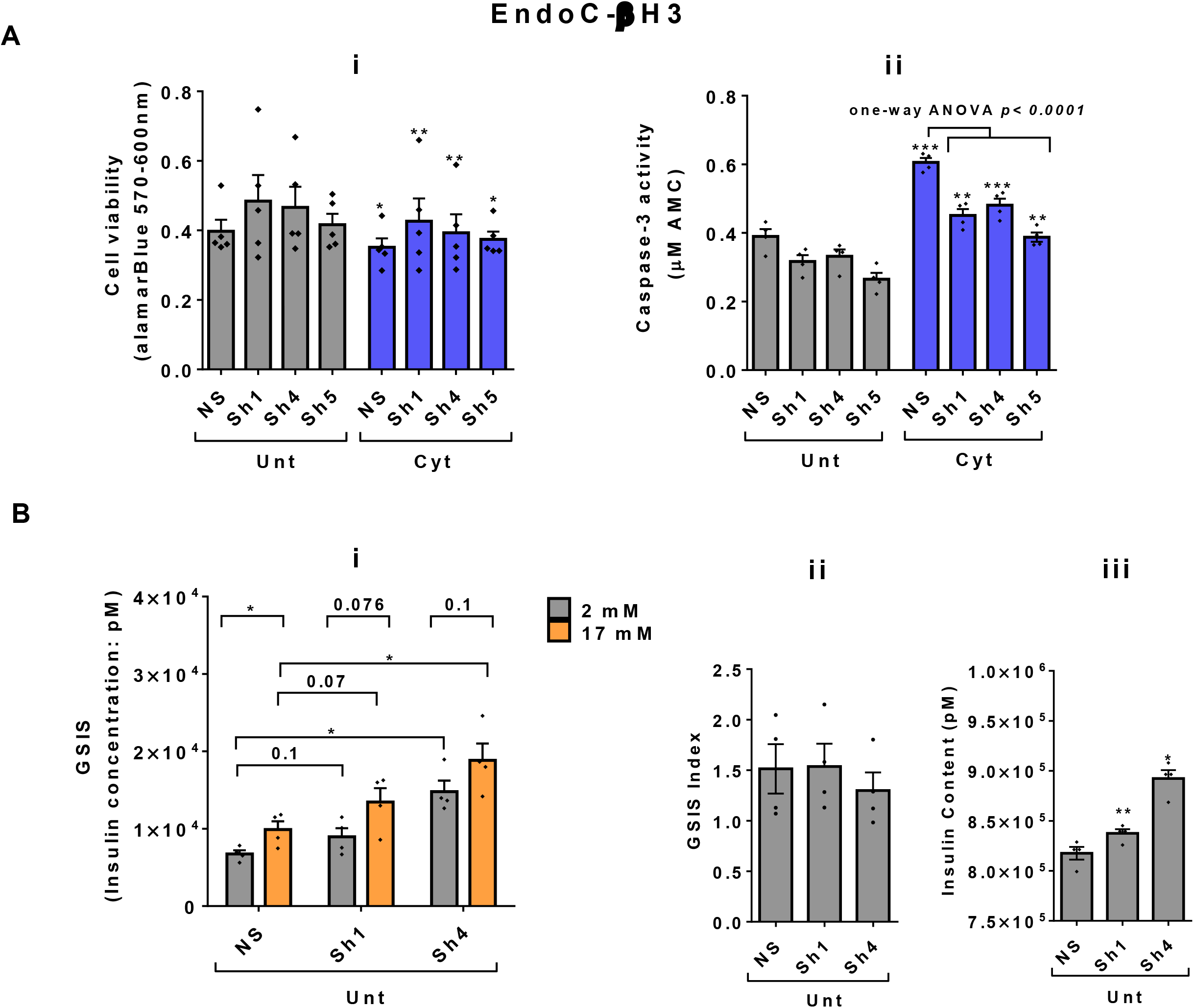
UPF2 knockdown potentiates cytokines suppression of NMD activity and slightly alleviates cytokines toxicity for cell viability and insulin content in EndoC-βH3 cells. EndoC-βH3 cell lines with the most efficient stable knock-down (KD) of UPF2 (three shRNAs; Sh1, Sh4 and Sh5) and non-silencing shRNA control (NS) were co-transfected with *Renilla*-HBB(PTC-) and or *Renilla*-HBB(PTC+), and the *Firefly* plasmids and exposed to PBS as untreated (Unt) and or cytokines combination (Cyt; 3 ng/mL IL-1β + 10 ng/mL IFNγ+10 ng/mL TNFα). A. Cell viability was measured by Alamarblue (A-i) and caspase-3 activity (A-ii) assays (N=6). B. Glucose-stimulated insulin secretion (GSIS) (B-i) and insulin contents (B-iii) were investigated in the UPF2 KD EndoC-βH3 cells. Insulin concentration (pM) was measured by human insulin ELISA (N=6). GSIS index (B-ii) was calculated by dividing insulin concentration measured in the treatments of 17 mM by 2 mM glucose. The data are means ± SEM. The symbol * indicates the Bonferroni-corrected paired t-test values of treatments versus untreated (Unt) NS control (A-B), otherwise, designated by a line on top of the bars or the Bonferroni-corrected paired t-test values of corresponding low versus high glucose, that is otherwise, designated by lines on the top of the bars (B-i). * ≤ 0.05, ** ≤ 0.01, *** ≤0.001.

These data indicate that the UPF2 plays a crucial role in cytokines-induced β-cell apoptosis. In addition, the increase in insulin content in UPF2 deficient EndoCβH3 cells implies that insulin transcripts could possibly be targets of the UPF2-dependent NMD pathway branch.

### UPF2 knockdown differentially affects cytokines- and glucolipotoxicity-mediated deregulation of EndoC-βH3 transcripts

Consistent with our observations above (Fig.5), we previously reported (11) that the deficiency of SMG6, an endoribonuclease and a key effector of NMD, rendered protection against cytokines-induced cell death and was associated with increased insulin content. Therefore, we aimed to identify potential NMD target transcripts by using RNA-sequencing to assess the transcriptome of cytokines- or PBS-treated EndoC-βH3 cells stably transfected with a non-silencing shRNA (NS) or the specific shRNA (shRNA-1 named U1) against *UPF2*. Due to the differential effects of Cyt and GLT on UPF3 expression cf. above, we also performed RNA-sequencing after UPF2 KD vs. NS control EndoC-βH3 exposed to GLT compared with PBS-treated.

The RNA-seq datasets from either UPF2 KD or NS control EndoC-βH3 cell lines exposed to cytokines and or GLT was dimensionally reduced by principal component analysis (PCA) into two main principal components, PC1 and PC2 (*p<0.05*). The PCA of the NS control EndoC βH3 cells demonstrated a high similarity between the biological replicates, a small within-group variance, and a distinct clustering of the untreated, cytokines and GLT groups (Supplementary Fig.7A-i and ii). Pearson correlation (*p<0.05*) between samples justified the clustering of biological replicates of cytokines, GLT and untreated conditions (Supplementary Fig.7B-i and ii). PCA revealed that the UPF2 knockdown increased the majority of variance in transcript isoforms of the cell, hence patterns leading to visually dispersed biological replicates from the cytokines-exposed isolates, whereas decreased the variances from the GLT and untreated biological replicates, hence clustered them together.

A Venn diagram of the RNA-seq datasets demonstrated around an approximate total of 14,000 commonly expressed transcripts (FDR<0.05) and those differentially expressed transcripts regulated by PBS (i.e., untreated control), cytokines or GLT (Supplementary Fig.7C-i and ii; Supplementary Fig.7D-i, ii, iii, and iv). Among them, cytokines regulated 54% (up) and 46% (down) (Fig.6A-i), whereas GLT impacted 48% (up) and 52% (down) of significantly expressed transcripts in NS control EndoC-βH3 cells (Fig.6A-iv). UPF2 KD changed the cytokines-mediated regulation of significantly expressed transcripts by 59% (up) and 41% (down) (Fig.6A-ii), whereas it did not change regulation of significantly expressed transcripts by GLT (up:47%, down:53%) in EndoC-βH3 cells (Fig.6A-v). This indicates UPF2 KD possibly alters the mRNA levels of identified transcript species.

**Figure 6.**
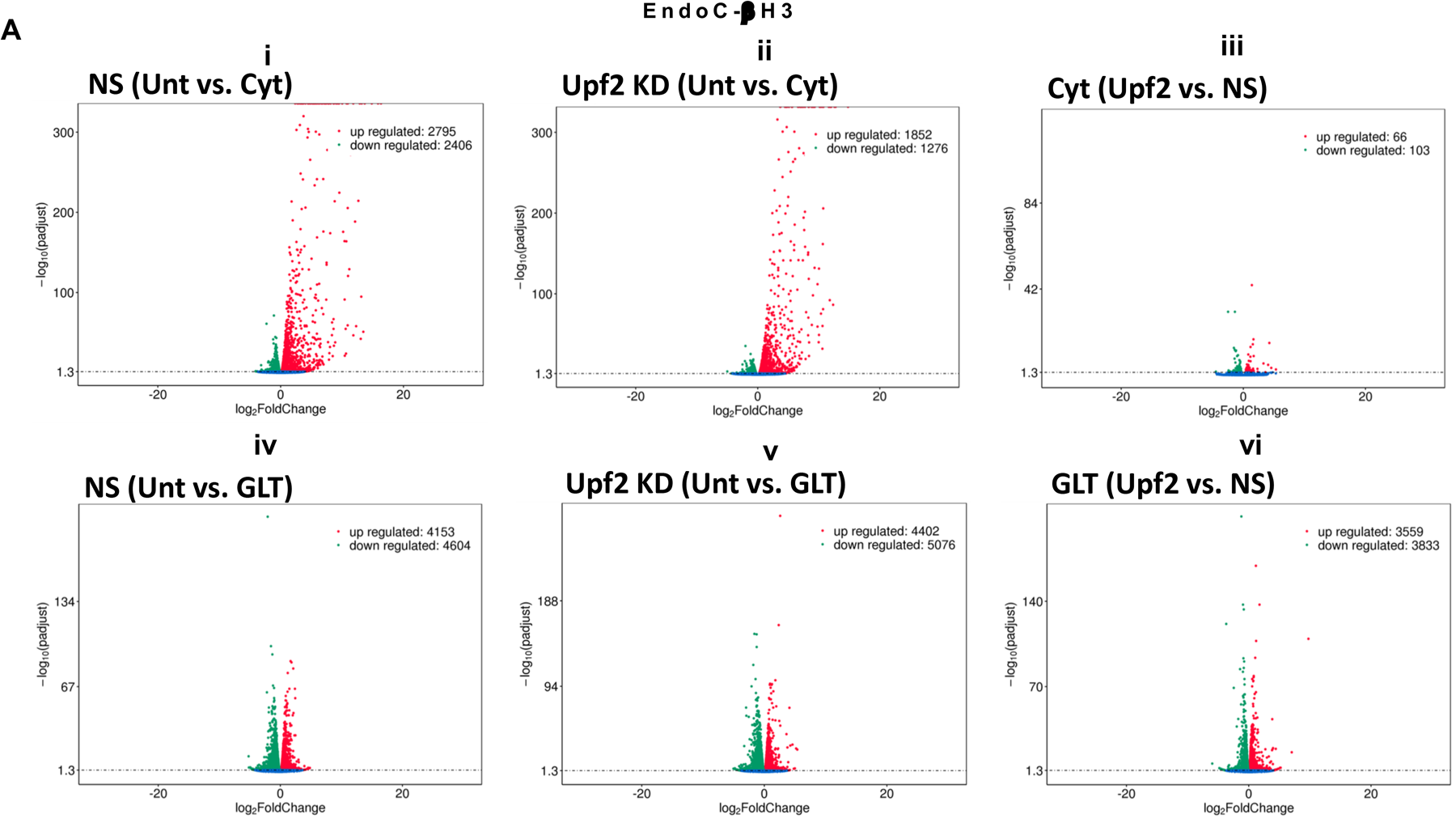
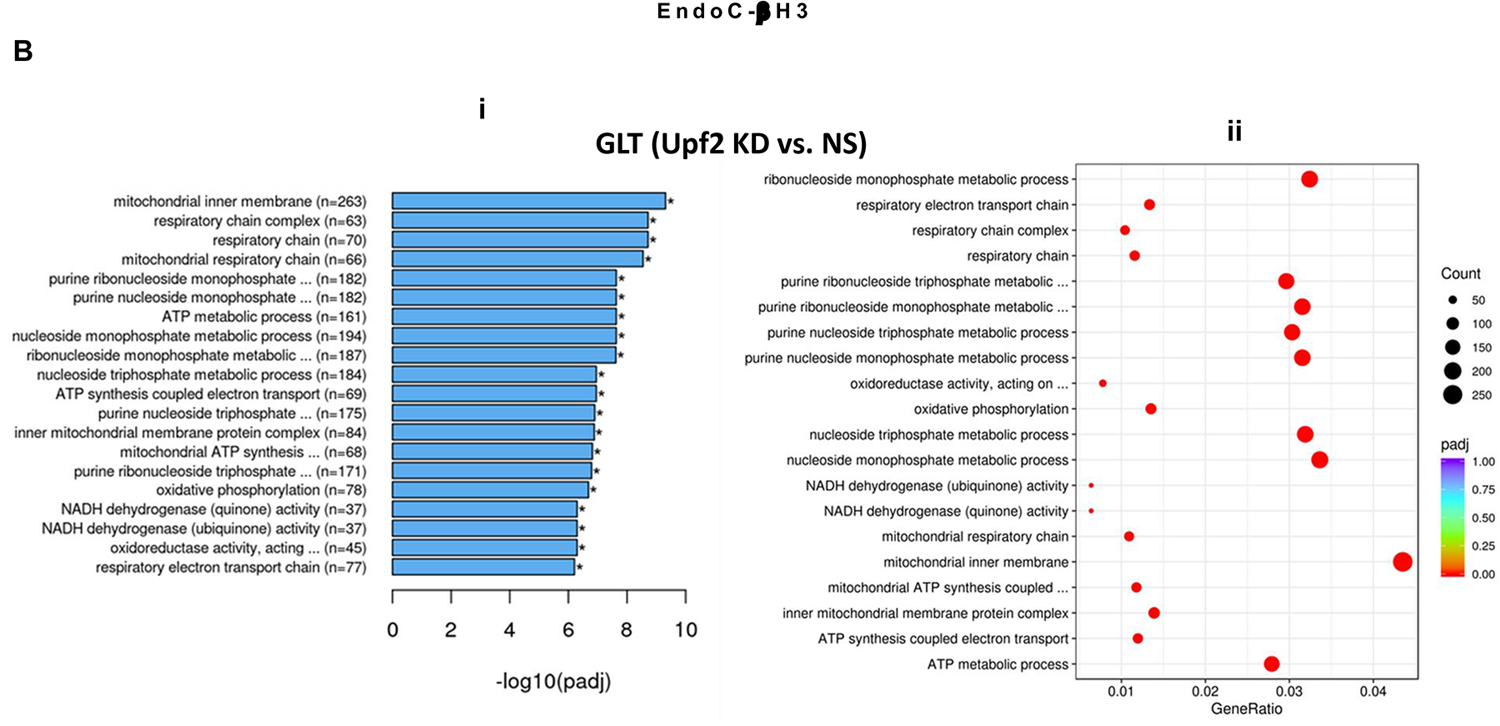
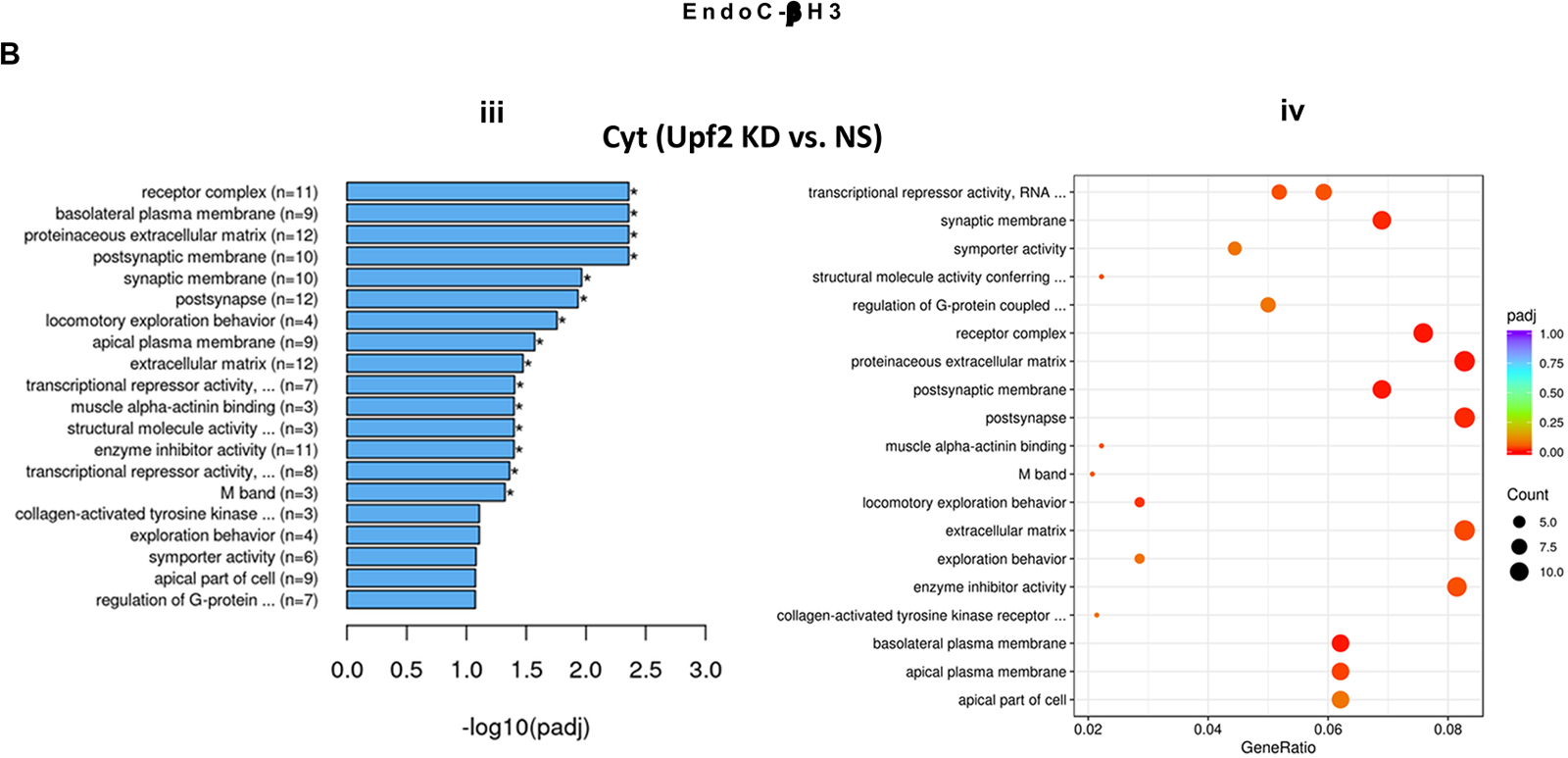
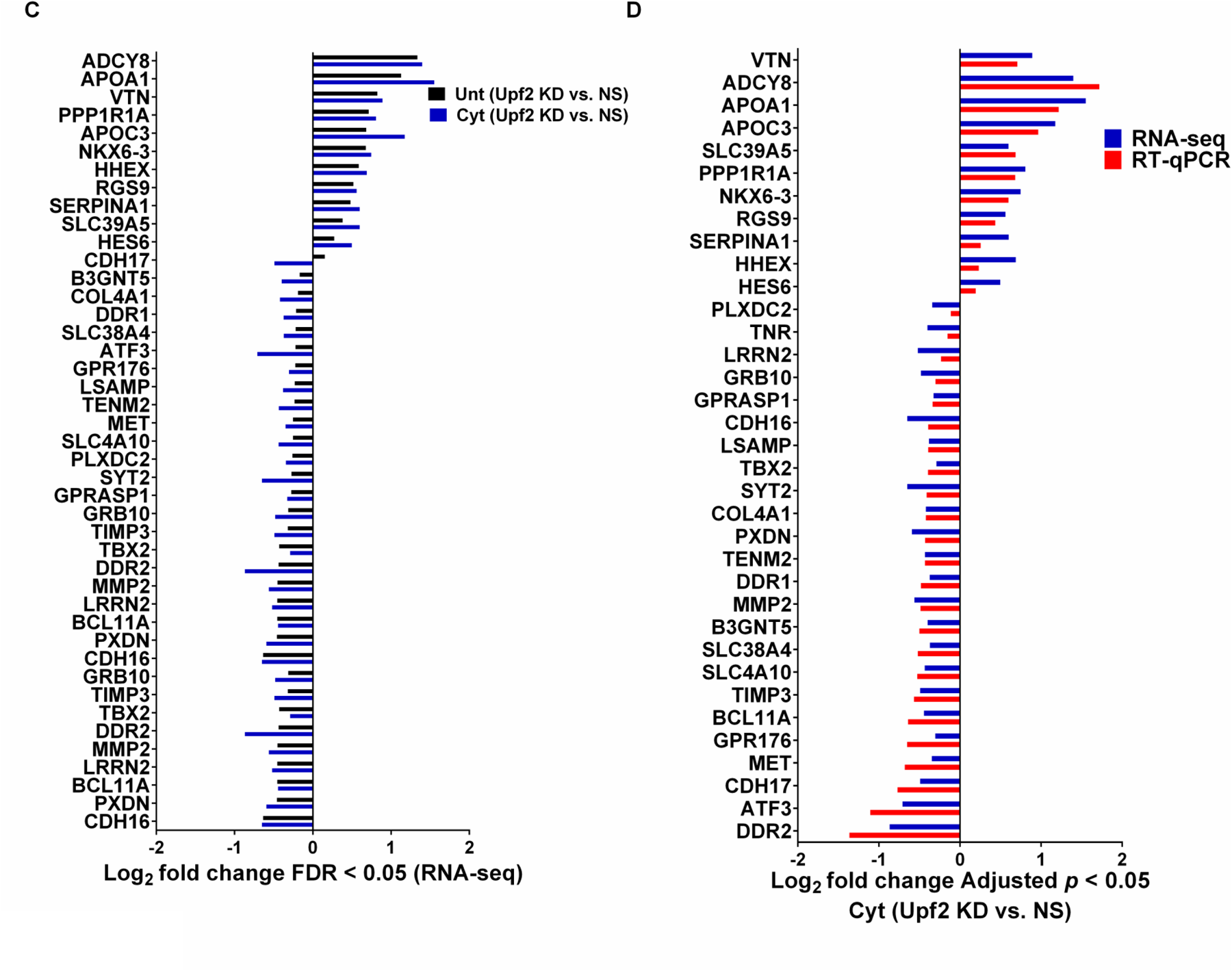
UPF2 knockdown differentially affects cytokines- and glucolipotoxicity-mediated deregulation of EndoC-βH3 transcripts. EndoC-βH3 cell lines knocked-down for UPF2 (shRNA-1 named U1) or non-silencing shRNA control (NS) were exposed to PBS as untreated (Unt), cytokines combination (Cyt; 3 ng/mL IL-1β + 10 ng/mL IFNγ+ 10 ng/mL TNFα) and or glucolipotoxicity (GLT; 0.5 mM Palmitate+25 mM glucose). Total RNA was extracted from the treated cells, cDNA library was made and sequenced using Hiseq platform as explained in methods. 33 RNA-seq datasets from NS/CTL (N=6), NS/Cyt (N=6), NS/GLT (N=4), U1/CTL (N=6), U1/Cyt (N=6) and U1/GLT (N=5) were analysed through the pipeline described in the supplementary methods. A. Volcano plot of number of transcripts (FDR <0.05) regulated by Cyt or GLT versus untreated in the NS or UPF2 KD (U1) cell lines. B. Top enriched pathways (subpanels i and iii) and the number of transcripts (subpanels ii and iv) regulated by Cyt and/or GLT in the UPF2 KD (FDR < 0.0.5). Enrichment is shown as log10 (adjusted *p*-value <0.05). C. Top GEA-identified transcripts regulated by Cyt compared to untreated in UPF2 KD vs. NS control EndoC-βH3 cells. The expression level is shown as log2 (adjusted *p*-value <0.05). D. RT-qPCR verification of the identified transcripts regulated by Cyt in UPF2 KD vs. NS control EndoC-βH3 cells. The expression level is shown as log2 (adjusted *p*-value <0.05).

To identify UPF2 KD-regulated transcripts possibly providing protection against cytokines-induced cytotoxicity, we interrogated cytokines-and/or GLT-regulated transcripts that were differentially expressed (*p<0.05*) in UPF2 KD versus NS control EndoC-βH3 cells using gene enrichment analysis (GEA). Gene Ontontolgy (GO) and Kyoto Encyclopaedia of Genes and Genomes (KEGG) gene enrichment analyses demonstrated that GLT significantly (*p<0.05*) regulated transcripts in the cellular functions of RNA splicing, mitochondrial inner membrane, and purine nucleoside metabolism (Supplementary Fig.7E-i and ii), although these were not affected by UPF2 KD (Fig.6B-i). In contrast, Gene Enrichment Analysis (GEA) revealed that in both untreated and cytokines-treated EndoC-βH3 cells, UPF2 KD significantly regulated transcripts encoding proteins involved in synaptic transmission, extracellular matrix, basolateral plasma membrane, receptor complex, synaptic membrane, transcriptional repressor activity and enzyme inhibitor activity (Fig.6B-iii; Fig.6C; Supplementary Fig.7E-iii and iv).

RT-qPCR confirmed the logarithmic fold change of UPF2 KD regulated transcripts in the cytokines-treated EndoC-βH3 cells *versus* cytokines-treated NS control cells (Fig.6D). The role of many of these transcripts in cell viability or insulin secretion was previously identified in pancreatic β-cells. Among them, α-1-antitrypsin (abbreviated 1-AT) has been proposed as an antagonist against cytokines-induced pancreatic β-cell death (27, 28). Our expression profiling verified that cytokines upregulated α-1-antitrypsin, and this effect was further potentiated by UPF2 KD. To explore the potential importance of these changes, we knocked down α-1-antitrypsin using specific siRNAs in INS1(832/13) and EndoC-βH3 cells as confirmed by quantitative WB (Supplementary Fig.8-i, ii and iii). The effect of siRNA-mediated α-1-antitrypsin knockdown was inconclusive in EndoC-βH3 cells (Supplementary Fig.7B-i and ii). Both α-1-antitrypsin siRNAs aggravated cytokine-induced cell death in comparison with NS control in INS1(832/13) cells (Fig.7A-iii and iv). Compared with NS control, α-1-antitrypsin knockdown reduced GSIS index, but had no effect on insulin content in INS1(832/13) cells (Fig.7B-ii, and iii), possibly due to the reduced cell number.

**Figure 7.**
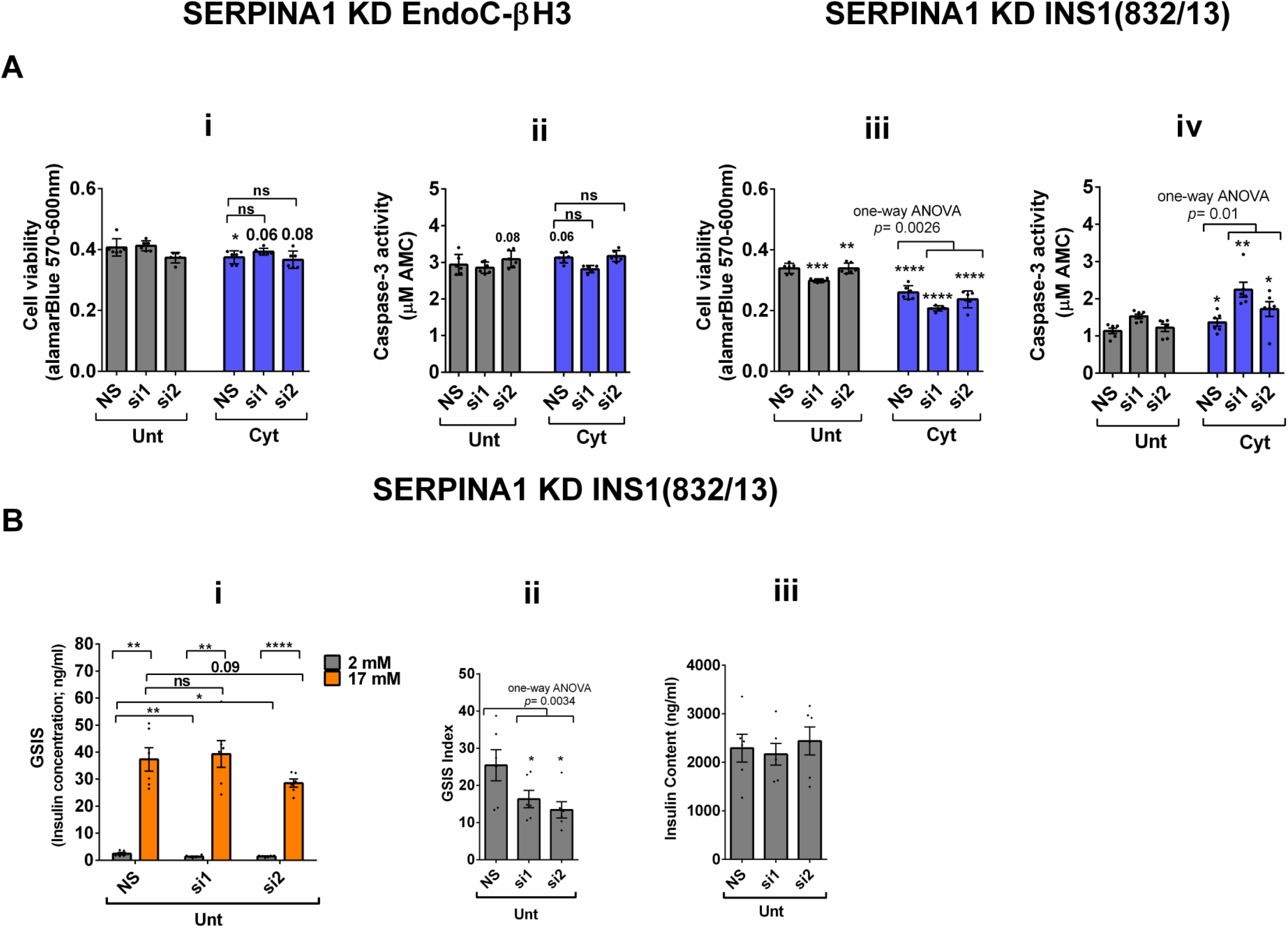
SERPINA1 knockdown deteriorates cytokines cytotoxicity for viability and glucose-stimulated insulin secretion index in INS1(832/13) cells. EndoC-βH3 and INS1(832/13) cells were transfected with siRNAs against SERPINA1 (two species-specific siRNAs for each cell type; si1 and si2) and a non-silencing siRNA control (NS), incubated for 24 hours and exposed to PBS as untreated (Unt) and cytokines combination (Cyt for EndoC-βH3; 3 ng/mL IL-1β + 10 ng/mL IFN-γ+ 10 ng/mL TNFα) (Cyt for INS1(832/13); 150 pg/mL IL-1β + 0.1 ng/mL IFNγ+ 0.1 ng/mL TNFα) for 72 and 24 hrs, respectively (see Supplementary Methods). The knockdown efficiency was checked using quantitative WB (Supplementary Fig.8-i, ii and iii). A. Cell viability was measured by Alamarblue (A-i and iii) and caspase-3 activity (A-ii and iv) assays (N=6). B. Glucose-stimulated insulin secretion (GSIS) (C-i) and insulin contents (C-iii) were investigated in the transfected EndoC-βH3 cells. Insulin concentration (ng/ml) was measured by insulin ultra-sensitive assay (N=6). GSIS index (C-ii) was calculated by dividing insulin concentration measured in the treatments of 17 mM by 2 mM glucose. The data are means ± SEM of N=6. The symbol * indicates the Bonferroni-corrected paired t-test values of treatments versus untreated (Unt) NS control or cytokines (Cyt)-treated NS that is, otherwise, designated by a line on top of the bars (A), or corresponding low versus high glucose that is, otherwise, designated by lines on the top of the bars (B-i). * ≤ 0.05, ** ≤ 0.01, *** ≤0.001, **** ≤ 0.0001. ns: non-significant.

Cytokines reportedly upregulate >30 splicing factors, affecting alternative splicing of 35% of genes in the human islet transcriptome (6). We examined RNA-seq datasets for alternative splicing (AS) isoforms (Supplementary Fig.9 i) driven by cytokines or GLT versus untreated in the NS control and UPF2 KD EndoC-βH3 cells. Among 2123 and 2106 cytokines-driven AS isoforms, skipped exon (SE) isoforms constituted 70.89% (*p*=0.1, n=6) and 72.5% (*p*=0.1, n=6) in NS control and UPF2 KD cells, respectively (Supplementary Fig.9-ii and iii). In contrast, 220 and 133 GLT-driven AS isoforms were identified in NS control and UPF2 KD, respectively (Supplementary Fig.9-iv and v). This differential regulation could possibly provide a reliable measure for cytokines and GLT role in inducing AS isoforms in β-cells.

Taken together, the above transcriptome analysis of EndoC-βH3 cells indicates that cytokines increase 1-AT expression, and this is synergised by NMD attenuation.

## Discussion

In this study, we demonstrate that cytokines decrease nonsense-mediated RNA decay (NMD) in INS1(832/13), EndoC-βH3 cells and dispersed human islets. We also showed that the cytokines-mediated decrease of NMD activity was driven by ER stress and downregulation of UPF3B. Loss-/or gain-of function of NMD activity could be elicited by UPF3B over-expression or UPF2 knockdown, which led to increases in, or slight decreases in, cytokines-induced apoptosis associated with decreased and increased insulin contents, respectively, without affecting GSIS index in EndoC-βH3 cells. Transcriptome profiling indicated a potentiating effect of UPF2 knockdown on Cyt, but not GLT-mediated, NMD activity. Interestingly, this approach identified transcript targets encoding proteins belonging to the extracellular matrix such as α-1-antitrypsin. Importantly, the knockdown of this gene enhanced cytokines-induced cytotoxicity in β-cells.

To the best of our knowledge the present study represents the first demonstration of a functional effect of cytokines on NMD activity.

UPR activation is known to inhibit NMD via PERK activation and eIF2 phosphorylation to restore IRE1α accumulation and hence a robust UPR activation (19, 20, 29); additional to the role of PERK activation, our findings suggest that IRE1α riboendonuclease activity (*p=0.1*) was involved in cytokines-mediated NMD inhibition in EndoC-βH3 cells.

UPF3A and UPF3B act as a potent NMD inhibitor and activator, respectively, in HeLa cells and in mice (23), consistent with our observations following UPF3B overexpression in β cells. However, the finding that forced UPF3A overexpression slightly increased NMD activity in β-cells seems inconsistent with previous findings. Recent studies (30, 31) support our apparently discrepant finding regarding the effects of UPF3A overexpression by showing redundancy of UPF3A and UPF3B as modular activators of NMD (24). With these two earlier studies in mind, we cannot rule out the interference of endogenous UPF3A in the actions of UPF3B on NMD in β-cells.

We investigated the consequences of NMD activity for pancreatic β-cell function and viability. The increase of NMD activity by UPF3B overexpression induced basal and cytokines-induced cell death in EndoC-βH3 cells, highlighting the role of increased UPF3B level in β-cells. This appears to be relevant for β-cell viability in both normal and inflammatory stress conditions. Similarly, UPF3B knockdown also caused basal cell death in INS1 cells. Hence, basal UPF3A/B levels seem to play crucial roles in the cell viability of β-cells and perturbation of such a controlled level implicates in cell death. On the other hand, the slight protection against cytokines-induced cell death conferred by UPF2 knockdown in EndoC-βH3 cells (Fig.5A-i and ii) and by SMG6 knockdown in INS1 cells (11) implies a possibly protective mechanism against cytotoxicity of cytokines in β-cells, irrespective to the outcome cytokines-induced cell death. Moreover, our findings provide evidence that the reduced insulin content observed after UPF3B overexpression is related to overactivated NMD.

Cytokines-induced perturbation of NMD, (potentiated by UPF2 silencing), might change the balance of anti-/pro-apoptotic transcripts. This, in turn, may contribute to cytotoxic damage. Consistent with this view, GEA revealed that cytokines deregulate transcripts encoding proteins that localise to and/or function in the extracellular matrix Thus, α-1-antitrypsin knockdown increased the detachment of MIN6 cells and exacerbated Thapsigargin-induced cell death as measured by Propidium-Iodide staining (28) and in this study increased cytokines-induced cell death in INS1(832/13) cells associated with decreased GSIS index.

We speculate that the perturbation of NMD by cytokines leads to increased exon skipping and that this may be part of a feedback loop promoting β-cell plasticity and resilience against cytotoxic cytokines. Future studies will be needed to test this possibility. We note also that depletion of alternative splicing factors (reviewed in (3)) inhibits insulin secretion and induces basal apoptosis and after treatment with cytokines in rodent and human β-cells (3, 7, 32, 33). Moreover, antisense-mediated exon skipping of 48–50 exons of the dystrophin gene restores the open reading frame and allows the generation of partially to largely functional protein (34).

In conclusion, we reveal that cytokines suppress NMD activity via ER stress signalling, possibly as a protective response against cytokines-induced NMD component expression. Our findings highlight the central importance of RNA turnover in β-cell responses to inflammatory stress.

### Limitations and Future Perspectives

We used a luciferase-based NMD reporter based on two separate PTC- and PTC+ constructs whose labelled luciferase is separately measured. Thus, a yet-to develop NMD activity reporter by which transcripts RNA, protein, or their corresponding labelled luciferase activity of both PTC- and PTC+ transcripts could be examined in one cell rather than (potentially) two separate cells will remove the limitation of current reporter based on transfection of the constructs into two separate cells. Moreover, the constant overexpression of UPF3A and UPF3B may result in cell death as β-cells cannot cope with overwhelming level of these proteins above the basal level. This could explain why UPF3B overexpression reduced the basal cell viability. Further studies should: 1) conduct in vivo experiments to validate the findings observed in vitro and determine if the regulation of NMD in pancreatic ß-cells is consistent across different cell types and conditions, 2) investigate the mechanisms underlying the regulation of NMD in pancreatic β-cells, including the role of specific NMD components and the impact of different stressors on NMD activity, 3) explore the potential therapeutic implications of targeting NMD in the treatment of inflammatory stress in β-cells, including the development of novel drugs or therapies that modulate NMD activity.

### Translatability of the findings

The findings report NMD involvement in the development of islet autoimmunity and the destruction of pancreatic β-cells in type 1 diabetes as well as islet inflammation in type 2 diabetes. The identification of novel targets arisen from cytokines-driven NMD attenuation could possibly suggest new biomarkers to monitor disease progression and may also guide the development of protein-based vaccines or antisense mRNA therapeutics in individuals who are at risk of diabetes development and or other inflammatory and autoimmune disorders.

## Supporting information

All supplemental data and figures

## Acknowledgments

Our special thanks to Dr. Gabriele Neu-Yilik, Professor Andreas E. Kulozik and Professor Matthias W. Hentze (Heidelberg University, Germany) for providing the plasmids *Renilla* (*RLuc)*-HBB PTC+, *RLuc*-HBB PTC-, *Firefly (FLuc)*, Upf3A, Upf3B and Upf3BΔ42 and our gratitude to Dr. Gabriele Neu-Yilik for initially reviewing with her comments and guidance on the NMD activity data. We thank Novogene (Cambridge, United Kingdom) for providing RNA-sequencing and RNA-seq raw data analysis. Human islets for research were provided by the Alberta Diabetes Institute Islet Core at the University of Alberta in Edmonton (http://www.bcell.org/adi-isletcore.html) with the assistance of the Human Organ Procurement and Exchange (HOPE) program, Trillium Gift of Life Network (TGLN), and other Canadian organ procurement organizations. Islet isolation was approved by the Human Research Ethics Board at the University of Alberta (Pro00013094). All donors’ families gave informed consent for the use of pancreatic tissue in research.

## Data accessibility

The RNA-seq data from the human insulin-producing cell line EndoC-βH3 that support the findings of Figure 6 of this study are deposited in the Sequence Read Archive (SRA) data repository (Accession numbers for 33 RNA-seq datasets: SRR22938756-SRR22938788) under the BioProject accession number (PRJNA916946) that are appreciated for further citations.

All other data generated or analysed during this study are included in this published article (and its supplementary information files).

## Author contributions

S.M.G. was lead investigator and grant holder (grant number: 9034-00001B) and principally conceptualized, designed, and performed all experiments, analysed data, prepared figures, and wrote the first draft. G.A.R. co-designed experiments, co-analysed data and co-wrote the manuscript. T.M.P. discussed scientific data and edited drafts. J.H.N. mentored the DFF fellowship application and edited the first draft. P.M. and L.P. provided human islets and discussed the data. B.P. discussed the UPF2 data and edited the first draft. All authors approved the manuscript.

## References

1. Rutter GA, Georgiadou E, Martinez-Sanchez A, Pullen TJ. Metabolic and functional specialisations of the pancreatic beta cell: gene disallowance, mitochondrial metabolism and intercellular connectivity. Diabetologia. 2020;63(10):1990–8.

2. Eizirik DL, Pasquali L, Cnop M. Pancreatic beta-cells in type 1 and type 2 diabetes mellitus: different pathways to failure. Nat Rev Endocrinol. 2020;16(7):349–62.

3. Alvelos MI, Juan-Mateu J, Colli ML, Turatsinze JV, Eizirik DL. When one becomes many-Alternative splicing in beta-cell function and failure. Diabetes Obes Metab. 2018;20 Suppl 2(Suppl 2):77–87.

4. Huang L, Wilkinson MF. Regulation of nonsense-mediated mRNA decay. Wiley Interdiscip Rev RNA. 2012;3(6):807–28.

5. Le K, Mitsouras K, Roy M, Wang Q, Xu Q, Nelson SF, et al. Detecting tissue-specific regulation of alternative splicing as a qualitative change in microarray data. Nucleic Acids Res. 2004;32(22):e180.

6. Eizirik DL, Sammeth M, Bouckenooghe T, Bottu G, Sisino G, Igoillo-Esteve M, et al. The human pancreatic islet transcriptome: expression of candidate genes for type 1 diabetes and the impact of pro-inflammatory cytokines. PLoS Genet. 2012;8(3):e1002552.

7. Ghiasi SM, Rutter GA. Consequences for Pancreatic beta-Cell Identity and Function of Unregulated Transcript Processing. Front Endocrinol (Lausanne). 2021;12:625235.

8. Hug N, Longman D, Caceres JF. Mechanism and regulation of the nonsense-mediated decay pathway. Nucleic Acids Res. 2016;44(4):1483–95.

9. Pan Q, Saltzman AL, Kim YK, Misquitta C, Shai O, Maquat LE, et al. Quantitative microarray profiling provides evidence against widespread coupling of alternative splicing with nonsense-mediated mRNA decay to control gene expression. Genes Dev. 2006;20(2):153–8.

10. Chang YF, Imam JS, Wilkinson MF. The nonsense-mediated decay RNA surveillance pathway. Annu Rev Biochem. 2007;76:51–74.

11. Ghiasi SM, Krogh N, Tyrberg B, Mandrup-Poulsen T. The No-Go and Nonsense-Mediated RNA Decay Pathways Are Regulated by Inflammatory Cytokines in Insulin-Producing Cells and Human Islets and Determine beta-Cell Insulin Biosynthesis and Survival. Diabetes. 2018;67(10):2019–37.

12. Asfari M, Janjic D, Meda P, Li G, Halban PA, Wollheim CB. Establishment of 2-mercaptoethanol-dependent differentiated insulin-secreting cell lines. Endocrinology. 1992;130(1):167–78.

13. Ravassard P, Hazhouz Y, Pechberty S, Bricout-Neveu E, Armanet M, Czernichow P, et al. A genetically engineered human pancreatic beta cell line exhibiting glucose-inducible insulin secretion. J Clin Invest. 2011;121(9):3589–97.

14. Boelz S, Neu-Yilik G, Gehring NH, Hentze MW, Kulozik AE. A chemiluminescence-based reporter system to monitor nonsense-mediated mRNA decay. Biochem Biophys Res Commun. 2006;349(1):186–91.

15. Neu-Yilik G, Raimondeau E, Eliseev B, Yeramala L, Amthor B, Deniaud A, et al. Dual function of UPF3B in early and late translation termination. EMBO J. 2017;36(20):2968–86.

16. Cunningham F, Allen JE, Allen J, Alvarez-Jarreta J, Amode MR, Armean IM, et al. Ensembl 2022. Nucleic Acids Res. 2022;50(D1):D988–D95.

17. Ghiasi SM, Hansen JB, Christensen DP, Tyrberg B, Mandrup-Poulsen T. The Connexin 43 Regulator Rotigaptide Reduces Cytokine-Induced Cell Death in Human Islets. Int J Mol Sci. 2020;21(12).

18. Vishnu N, Hamilton A, Bagge A, Wernersson A, Cowan E, Barnard H, et al. Mitochondrial clearance of calcium facilitated by MICU2 controls insulin secretion. Mol Metab. 2021;51:101239.

19. Karam R, Lou CH, Kroeger H, Huang L, Lin JH, Wilkinson MF. The unfolded protein response is shaped by the NMD pathway. EMBO Rep. 2015;16(5):599–609.

20. Goetz AE, Wilkinson M. Stress and the nonsense-mediated RNA decay pathway. Cell Mol Life Sci. 2017;74(19):3509–31.

21. Csutora P, Su Z, Kim HY, Bugrim A, Cunningham KW, Nuccitelli R, et al. Calcium influx factor is synthesized by yeast and mammalian cells depleted of organellar calcium stores. Proc Natl Acad Sci U S A. 1999;96(1):121–6.

22. Sehgal P, Szalai P, Olesen C, Praetorius HA, Nissen P, Christensen SB, et al. Inhibition of the sarco/endoplasmic reticulum (ER) Ca(2+)-ATPase by thapsigargin analogs induces cell death via ER Ca(2+) depletion and the unfolded protein response. J Biol Chem. 2017;292(48):19656–73.

23. Shum EY, Jones SH, Shao A, Dumdie J, Krause MD, Chan WK, et al. The Antagonistic Gene Paralogs Upf3a and Upf3b Govern Nonsense-Mediated RNA Decay. Cell. 2016;165(2):382–95.

24. Yi Z, Arvola RM, Myers S, Dilsavor CN, Alhasan RA, Carter BN, et al. Mammalian UPF3A and UPF3B activate NMD independently of their EJC binding. bioRxiv. 2021:2021.07.02.450872.

25. Serdar LD, Whiteside DL, Baker KE. ATP hydrolysis by UPF1 is required for efficient translation termination at premature stop codons. Nat Commun. 2016;7:14021.

26. Neu-Yilik G, Gehring NH, Hentze MW, Kulozik AE. Nonsense-mediated mRNA decay: from vacuum cleaner to Swiss army knife. Genome Biol. 2004;5(4):218.

27. Lewis EC, Mizrahi M, Toledano M, Defelice N, Wright JL, Churg A, et al. alpha1-Antitrypsin monotherapy induces immune tolerance during islet allograft transplantation in mice. Proc Natl Acad Sci U S A. 2008;105(42):16236–41.

28. McKimpson WM, Chen Y, Irving JA, Zheng M, Weinberger J, Tan WLW, et al. Conversion of the death inhibitor ARC to a killer activates pancreatic beta cell death in diabetes. Dev Cell. 2021;56(6):747–60 e6.

29. Karousis ED, Nasif S, Muhlemann O. Nonsense-mediated mRNA decay: novel mechanistic insights and biological impact. Wiley Interdiscip Rev RNA. 2016;7(5):661–82.

30. Wallmeroth D, Boehm V, Lackmann J-W, Altmüller J, Dieterich C, Gehring NH. UPF3A and UPF3B are redundant and modular activators of nonsense-mediated mRNA decay in human cells. bioRxiv. 2021:2021.07.07.451444.

31. Wallmeroth D, Lackmann JW, Kueckelmann S, Altmuller J, Dieterich C, Boehm V, et al. Human UPF3A and UPF3B enable fault-tolerant activation of nonsense-mediated mRNA decay. EMBO J. 2022;41(10):e109191.

32. Villate O, Turatsinze JV, Mascali LG, Grieco FA, Nogueira TC, Cunha DA, et al. Nova1 is a master regulator of alternative splicing in pancreatic beta cells. Nucleic Acids Res. 2014;42(18):11818–30.

33. Juan-Mateu J, Rech TH, Villate O, Lizarraga-Mollinedo E, Wendt A, Turatsinze JV, et al. Neuron-enriched RNA-binding Proteins Regulate Pancreatic Beta Cell Function and Survival. J Biol Chem. 2017;292(8):3466–80.

34. Aartsma-Rus A, van Ommen GJ. Antisense-mediated exon skipping: a versatile tool with therapeutic and research applications. RNA. 2007;13(10):1609–24.

