## Supplementary material for "Proinflammatory Cytokines Suppress Nonsense-Mediated RNA Decay to Impair Regulated Transcript Isoform Processing in Pancreatic β-Cells": All supplemental data and figures

and Guy A. Rutter^1,5,6*^

1. Section of Cell Biology and Functional Genomics, Division of Diabetes, Endocrinology and Metabolism, Department of Metabolism, Digestion and Reproduction, Faculty of Medicine, Imperial College London Du Cane Road, London W12 0NN, United Kingdom
2. Department of Biomedical Sciences, University of Copenhagen, 3 Blegdamsvej, 2200 Copenhagen N, Denmark.
3. Department of Clinical and Experimental Medicine, Islet Cell Laboratory, University of Pisa, 56126, Pisa, Italy
4. Diabetes Research Institute, IRCCS Ospedale San Raffaele, Milano, Italy
5. CHUM Research Centre (CRCHUM), University of Montreal, 900 Rue St. Denis, Montreal, QC, Canada
6. Lee Kong Chian School of Medicine, Nanyang Technological University, 637553, Singapore
7. Biotech Research and Innovation Centre (BRIC), University of Copenhagen, 3 Blegdamsvej, 2200 Copenhagen N, Denmark
8. The Finsen Laboratory, Rigshospitalet, Faculty of Health Sciences, University of Copenhagen, Copenhagen, Denmark
9. Danish Stem Cell Center (DanStem) Faculty of Health Sciences, University of Copenhagen, Copenhagen, Denmark
10. Sanford Burnham Prebys Medical Discovery Institute, La Jolla, CA, USA

**^†^** Lead investigator: m

### Supplementary Materials and Methods

**Materials and Preparations**

Recombinant rat and human IL-1β, IFNγ, and rhTNFα were purchased from R&D Systems (Human cytokines: Cat#201-LB-025/CF, 285-IF-100/CF. 210-TA-100/CF, Rat cytokines: Cat# 501-RL-010/CF, 585-IF-100/CF, 510-RT-050/CF, Minneapolis, USA) and tamoxifen from Sigma (Cat# T5648-5G, London, England).

Complete culture media specific for the rat or human β-cell lines and human islets:

For INS-1(832/13) cell pre-culture and experiments RPMI-Glutamax with 11 mM glucose (Cat#61870-010, LifeTechnologies), 10% FBS (Cat# A3840002, LifeTechnologies, Renfrew, England), 1% Penicillin/Streptomycin (P/S; Cat#15140122, LifeTechnologies) and 50 µM 2-mercaptoethanol (Cat#M6250, Sigma).

For human islet preculture RPMI (Cat#11879-020, LifeTechnologies) supplemented with 5.6 mM glucose (Cat#50-152-108, LifeTechnologies), 10% heat-inactivated FBS (Cat#10082147, LifeTechnologies), and 1% P/S; for human islet experiments RPMI (Cat#11879-020, LifeTechnologies) supplemented with 5.6 mM glucose, 2% human serum (Cat#11966025, Cambridge Bioscience; Cambridge, England), and 1% P/S.

For human insulin-producing EndoC-βH3 cells: DMEM (cat #11966025, LifeTechnologies) supplemented with 5.6 mM glucose, 2% BSA-fraction V, 50 μM 2-mercaptoethanol, 10 mM nicotinamide (Cat#N3376, Sigma), 5.5 mg/ml human transferrin (Cat#T3309, Sigma), 6.7 ng/ml sodium selenite (Cat#S5261, Sigma), Penicillin/Streptomycin (Cat#15140122, LifeTechnologies) and Puromycin (Cat#A1113803, LifeTechnologies) (1).

Glucolipotoxic conditions (GLT) were created with 0.5 mM palmitate (cat # P9767, Sigma) conjugated with 0.1% BSA-fraction V (Cat#10735078001, Sigma) + 25 mM glucose as described in (2, 3).

**Cell culture, human islet dispersion and treatment**

The human EndoC-βH3 and rat INS1(832/13) cell lines were tested negative for *Mycoplasma* and maintained as previously described (1, 4). EndoC-βH3 cells were matured with 1µM of tamoxifen (TAM) for three weeks according to the protocol previously described (1). The TAM-treated EndoC-βH3 cells were seeded onto fibronectin-and ECM-coated culture plates for further experiments (1).

Islets from eight human heart-beating organ-donors (Supplementary Table 1) (>80% purity, donor characteristics listed in Suppl. Table 1A) were isolated under local ethical approval, received in fully anonymous form and pre-cultured as described previously (5). There were no apparent differences in results obtained with islets from male or female donors, and data were therefore combined. For dispersion, 1000 islets were picked after 3 days of pre-incubation, spun down at 1000 rpm for 1 min, washed three times with PBS buffer (w/o Ca^2+^ or Mg^2+^, Cat# 14190250, Life Technologies), trypsinised with 500 µl Trypsin-EDTA 0,05% (Cat# 25200056; LifeTechnologies) and incubated at 37^o^C for 3 min. The digested islets were dispersed into single cell suspension by one minute of pipetting. Then, 15 ml complete medium was added to neutralize trypsin, and the dispersed cells were centrifuged at 1500 rpm for 5 min, re-suspended in the complete medium, mixed by transfection cocktail, re-plated in Fibronectin/ECM-coated 48-well plates and incubated for 48 h.

INS1(832/13), EndoC-βH3 or dispersed human islet cells were seeded in either 6-well plates (1×10^6^ cells/well for RNA isolation), 48-well plates (50,000 cells /well for luciferase assay) or 96-well plates (30,000 cells/well for Alamarblue and Caspase-3 activity assays), respectively. After 48h of pre-incubation, INS1 (832/13), EndoC-βH3 or dispersed human islet cells were exposed to cytokines mixture (Cyt) at the concentrations indicated in figures or figure legends for the indicated time points, glucolipotoxic (GLT) conditions, or PBS as negative control.

**Luciferase activity assay and NMD activity**

Plasmids (650 ng) encoding either human haemoglobin-β (HBB): HBB(NS39 or namely PTC+) or HBB(wildtype; WT or namely PTC-) fused with *Renilla (RLuc)*, and *Firefly (FLuc)* plasmid (PMID: 16934750) as transfection efficiency reference were mixed with 250 µl OptiMEM (Cat#31985062, LifeTechnologies), then with 20 µl FUGENE (Cat#E2311, Promega, Southampton, England), vortexed and incubated for 30 minutes at room temperature (RT). Monolayer INS1 (832/13) and EndoC-βH3 cultures were trypsinised into single cells and washed twice with Ca^2+^/Mg^2+^-containing PBS buffer (Cat#14080055, LifeTechnologies), whereafter the cell pellets were re-suspended in 2 ml complete medium. One million of either the split INS1(832/13) or EndoC-βH3 cells were added to the transfection cocktail and mixed by pipetting. The transfected cells were seeded in 48-well plates (1×10^5^ cells /well) and pre-incubated for 48 h in 37 ºC with 5% CO2.

*Renilla* and *Firefly* luminescence (*RLuc* and *FLuc*) was measured in a Lumat LB 9507 Tube Luminometer (Berthold Technologies, Bad Wildbad, Germany) by Dual-Luciferase Reporter Assay (Cat# E1910, Promega, Hampshire, England) according to the manufacturer’s manual, after lysis of a volume of 20 µl of the transfected cells by adding 100 µl of Passive Lysis Buffer. Luciferase Assay Substrate and Stop&Glo Buffer were manually injected immediately and mixed by 4 times pipetting. The integration time was set on the machine to 1 s after a 4 s delay time. *Renilla (RLuc)* luminescence signals were normalized to the *Firefly (FLuc)* control in both HBB(PTC+) and HBB(PTC-) (6). NMD activity was calculated by dividing the *RLuc/FLuc*-HBB(PTC-) by the *RLuc/FLuc*-HBB (PTC+) as illustrated in supplementary Fig. 1A and originally described (6). Experiments where the control *RLuc/FLuc-*HBB(PTC-) was affected by Cyt were excluded, so that the resulting NMD activity only denotes the PTC-containing HBB(PTC+).

**Functional analysis of UPF3A/B overexpression**

Expression plasmids (650 ng) encoding UPF3A, UPF3B or UPF3BΔ42 (7) were mixed with 500 µl OptiMEM (Cat#31985062, LifeTechnologies), then with 40 µl FUGENE (Cat# E2311, Promega, Southampton, England), vortexed and incubated for 30 minutes at room temperature (RT). Within the incubation time, the INS1(832/13) and EndoC-βH3 single cells were prepared as above, and one million cells were added to respective transfection cocktail and mixed by pipetting. The transfected cells were re-counted and seeded in 6-well plates (0.5×10^6^ cells/well for Western blot analysis of overexpression), 12-well plates (0.3×10^6^ cells/well for GSIS) or 96-well plates (0.03×10^6^ cells/well for Alamarblue and Caspase 3 activity assays), respectively, and pre-incubated for 48 h at 37ºC with 5% CO_2_.

To examine the impact of UPF3A/B overexpression on NMD activity, the transfection cocktail prepared above was divided into two, then 325 ng plasmid of either HBB(PTC+) or HBB(PTC-), and with the *Firefly* plasmid were added, followed by vortexing and incubation for 30 min at RT. Five hundred cells of the cell-splits were added as above to the transfection cocktail, mixed by pipetting, seeded in 48-well plates (1×10^5^ cells /well) and pre-incubated for 48 hours in 37ºC with 5% CO_2_.

**Lentiviral shRNA particle production**

GPIZ lentiviral shRNA Kits containing shRNA vectors for *Upf2, Upf3A* and *Upf3B* genes (Supplementary Table 3) along with a non-silencing shRNA (NS) vector as negative control were from Horizon Discovery (Cambridge, United Kingdom). All plasmids contained a puromycin resistance gene. HEK293FT cells were pre-incubated with OPTI-MEM medium (Cat#11058021, LifeTechnologies) without antibiotics 2h prior to transfection. Lentiviral shRNA particles were produced using the Trans-Lentiviral shRNA Packaging System (Cat#TLP5914, Horizon Discovery, Cambridge, England) according to the manufacturer’s instructions. The lentiviral particles were concentrated using Lenti-X™ Concentrator (Cat#631231, Takara, California, USA) according to the manufacturer’s instructions. The concentrated virus pellet was dissolved in 1 ml culture medium and stored at -80°C. Multiplicity of infection (MOI) was determined in INS1(832/13) cells using a 10-fold serial dilution along with puromycin-resistance selection (2 µg/ml). After 48h, the MOI of each shRNA lentivirus was scored by counting surviving cells under an inverted fluorescence microscope. Transduced INS1(832/13) and EndoC-βH3 cells were sorted by FACS and reseeded to reach a clonal expansion. The efficiency of knockdown was examined by Western blot analysis.

**Library preparation for transcriptome sequencing**

Total RNA from the NS control and or UPF2 KD EndoC-βH3 cells treated with cytokines, GLT, or PBS was extracted using TRIZOL (Cat#15596018, LifeTechnologies) with DNase (Cat#EN0521, Life Technologies) treatment and isopropanol precipitation according to the manufacturer’s instructions. Quality and quantity of the extracted total RNA from 33 independent biological replicates (i.e., N=6 of each PBS-exposed and or cytokines-exposed NS control and UPF2 KD, and N=4/N=5 of GLT-exposed NS control/UPF2 KD, respectively) were assessed using a NanoDrop-1000 (Thermo Fisher Scientific, Dartford, England). Total RNA (1 µg per isolate) was used as input material for the RNA sample preparations. Sequencing libraries were generated using NEBNext® Ultra TM RNA Library Prep Kit for Illumina® (NEB, Ipswich, USA) following manufacturer’s recommendations.

Briefly, mRNA was purified from total RNA using poly-T oligo-attached magnetic beads. Fragmentation was carried out using divalent cations under elevated temperature in NEBNext First Strand Synthesis Reaction Buffer (5X). First strand cDNA was synthesized using random hexamer primer and M-MuLV Reverse Transcriptase (RNase H). Second strand cDNA synthesis was subsequently performed using DNA Polymerase I and RNase H. Remaining overhangs were converted into blunt ends via exonuclease/polymerase activities. After adenylation of 3’ ends of DNA fragments, NEBNext Adaptor with hairpin loop structure were ligated to prepare for hybridization. In order to select cDNA fragments of preferentially 150~200 bp in length, the library fragments were purified with AMPure XP system (Beckman Coulter, Beverly, USA). Then 3 µl USER Enzyme (NEB) was used with size-selected, adaptor-ligated cDNA at 37 °C for 15 min followed by 5 min at 95 °C before PCR. Subsequently, PCR was performed with Phusion High-Fidelity DNA polymerase, Universal PCR primers and Index (X) Primer. Finally, PCR products were purified using AMPure XP, and library quality was assessed on the Agilent Bioanalyzer 2100 system (Agilent Technologies, California, USA).

Index codes were added to attribute sequences to each sample. The clustering of the index-coded samples was performed on a cBot Cluster Generation System using PE Cluster Kit cBot-HS (Illumina, San Diego, USA) according to the manufacturer’s instructions. After cluster generation, the library preparations were sequenced on an Illumina platform and paired-end reads were generated.

**Transcriptomic data analysis**

Raw data (raw reads) of FASTQ format were firstly processed through *fastp*. In this step, clean data (clean reads) were obtained by removing reads containing adapter and poly-N sequences and reads with low quality from raw data. All the downstream analyses were based on the clean data with high quality. The RNA-seq clean data were mapped to human reference genome (<http://www.ensembl.org/Homo_sapiens/Info/Index>) using the Spliced Transcripts Alignment to a Reference (STAR) software. The FeatureCounts algorithm was used to count the read numbers mapped to each gene, then Reads Per Kilobase of exon model per million mapped reads (RPKM) of each gene were calculated based on the length of the gene, and reads count mapped to this gene.

Principal component analysis (PCA), which was used to classify expression patterns according to gene expression level variance, was performed as previously described (8). PCA geometrically projects the high-dimensional dataset onto lower dimensions called principal components (PCs), with the goal of finding the best summary of the data, using a limited number of PCs. The first PC (i.e., PC1) was chosen to minimize the total distance between the data and their projection onto the PC. The second PC (i.e., PC2) was selected similarly, with the additional requirement that it was uncorrelated with PC1(8).

Differential expression analysis *between conditions/groups* (i.e., untreated and cytokines or GLT, or NS control and UPF2 KD (4-6 biological replicates per condition)) was performed using the DESeq2 R package. DESeq2 provides statistical routines for determining differential expression in digital gene expression data using a model based on negative binomial distribution. The resulting *p* values were adjusted using the Benjamini and Hochberg’s approach for controlling the False Discovery Rate (FDR). Genes with an adjusted *P* value < 0.05 found by DESeq2 were assigned as differentially expressed.

Differential expression analysis of *two conditions* was performed using the *edgeR* R package. The *P* values were adjusted using the Benjamini and Hochberg method. Corrected *P* value of 0.005 and |log2 (Fold Change) | of 1 were set as the threshold for significantly differential expression. Gene enrichment analysis (GEA) was performed by biological ontologies including Gene Ontology (GO), Kyoto Encyclopaedia of Genes and Genomes (KEGG), Reactome, Human Disease Ontology (DO) and DisGeNET (<https://www.disgenet.org>). The *clusterProfiler*, an R package, was used for statistical enrichment of differentially expressed genes, and the gene ontology terms with adjusted *P* value < 0.05 were considered significant enrichment.

Alternative splicing analysis was performed by the software *rMATS*, that identifies alternative splicing events corresponding to all major types of alternative splicing patterns and calculates the *P* value and FDR for differential splicing. These types include skipped exons (SE), alternative 5’ splice sites (5SS), alternative 3’ splice sites (3SS), mutually exclusive exons (MXE), and intron retentions (IR).

**cDNA synthesis and RT-qPCR**

Purified total RNA (500 ng) was used for cDNA synthesis with the SuperScript™ Kit (Cat#18090010, LifeTechnologies). Real-time Reverse Transcriptase-quantitative PCR (RT-qPCR) was performed on 12 ng cDNA with SybrGreen PCR mastermix (Life Technologies) and specific primers (Supplementary Table 2) and run in a real-time PCR machine (Applied Biosystems QuantStudio, Thermo Fisher Scientific). NormFinder software (9) was used to select the most stable reference gene among Hprt1, Actin and Tubulin. Statistical analysis was carried out on the gene expression levels normalized to the chosen reference gene (see figure legends) through -∆Ct analysis, and figures are presented as logarithmic fold change versus Ctl which was calculated by the -∆∆Ct method.

**Western blotting (WB) analysis**

Cells were lysed on ice with NP-40 lysis buffer containing protease inhibitor cocktail (LifeTechnologies) and stored at -20^o^C. Five µg of lysate from each condition of the same experiment adjusted for protein concentration with Bradford assay according to the manufacturer’s protocol (Bio-Rad) were loaded in duplicate-quadruplicate on the same gel and were separated by 4-20% SDS-PAGE as appropriate and blotted on PDVF membrane (Bio-Rad). To avoid stripping membranes when staining for multiple proteins, membranes were cut according to the desired molecular weight (MW) range, stained with antibodies against alpha-Tubulin (1:2000) (Cat#T5168, Sigma), UPF2 (1:1000) (LifeTechnologies), UPF3A (1:1000) (Cat#PA5-41904, LifeTechnologies), UPF3B (1:1000) (Cat#PB9843, Boster Bio, Pleasanton, USA) and α-1-antitrypsin (1:1000) (Cat#TA500375, LifeTechnologies), and developed with the chemiluminescence detection system Super Signal (Life Technologies) as previously described (10). High-quality blots of the replicates were quantified relative to the respective loading Tubulin controls. Light emission was captured using an Alphaimager system (Alpha-Innotech, Exeter, England). Band density was quantified using ImageJ software.

### Supplementary tables

### Table S1. List of human islet donor characteristics

| **No.** | **Provider** | **Sex** | **Age** | **BMI** |
| --- | --- | --- | --- | --- |
| **1** | **Pisa** | Male | 63 | 29.4 |
| **2** | **Edmonton** | Male | 9 | 18.1 |
| **3** | **Pisa** | Female | 64 | 22.07 |
| **4** | **Milan** | Female | 53 | 27.2 |
| **5** | **Edmonton** | Male | 42 | 29.9 |
| **6** | **Edmonton** | Male | 41 | 33.1 |
| **7** | **Pisa** | Female | 76 | 23.9 |
| **8** | **Pisa** | Male | 80 | 23.03 |

### Table S2. List of specific primers for RT-qPCR. (* taken from (10))

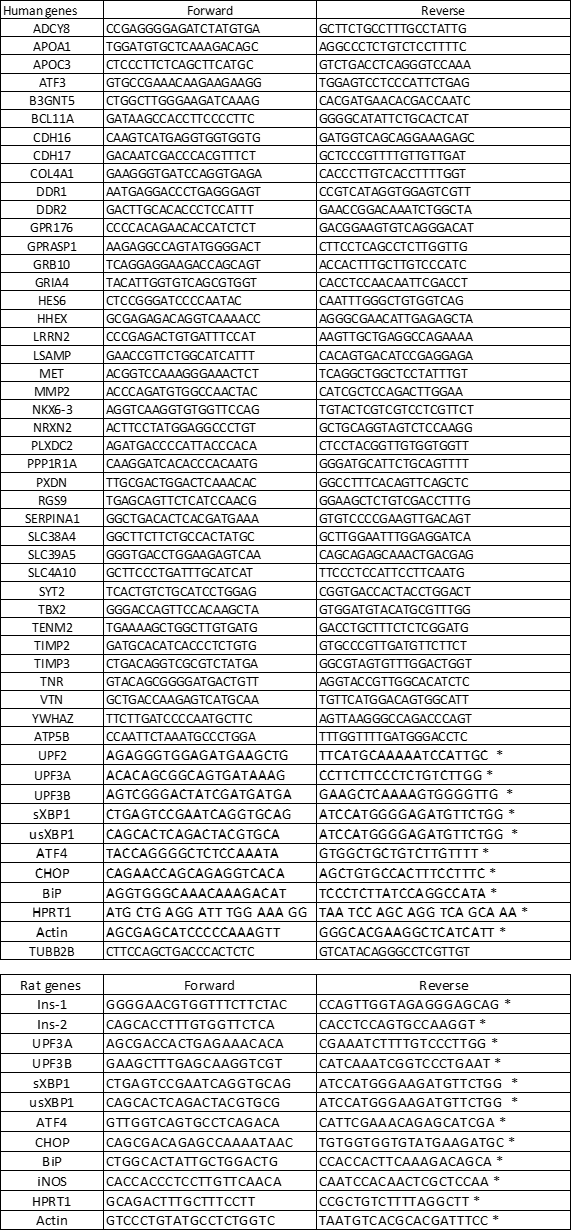

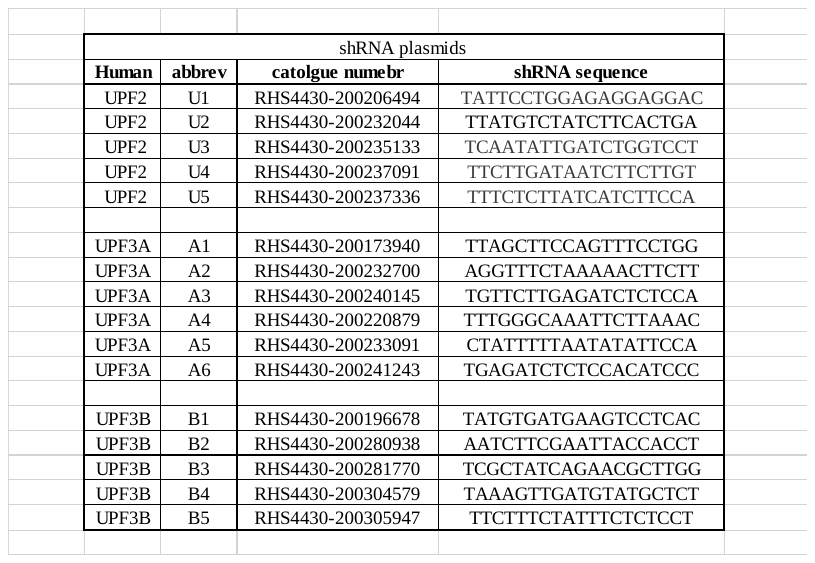

### Table S3. List of gene-specific shRNA sequences.

### Supplementary Figures

### Supplementary Figure 1

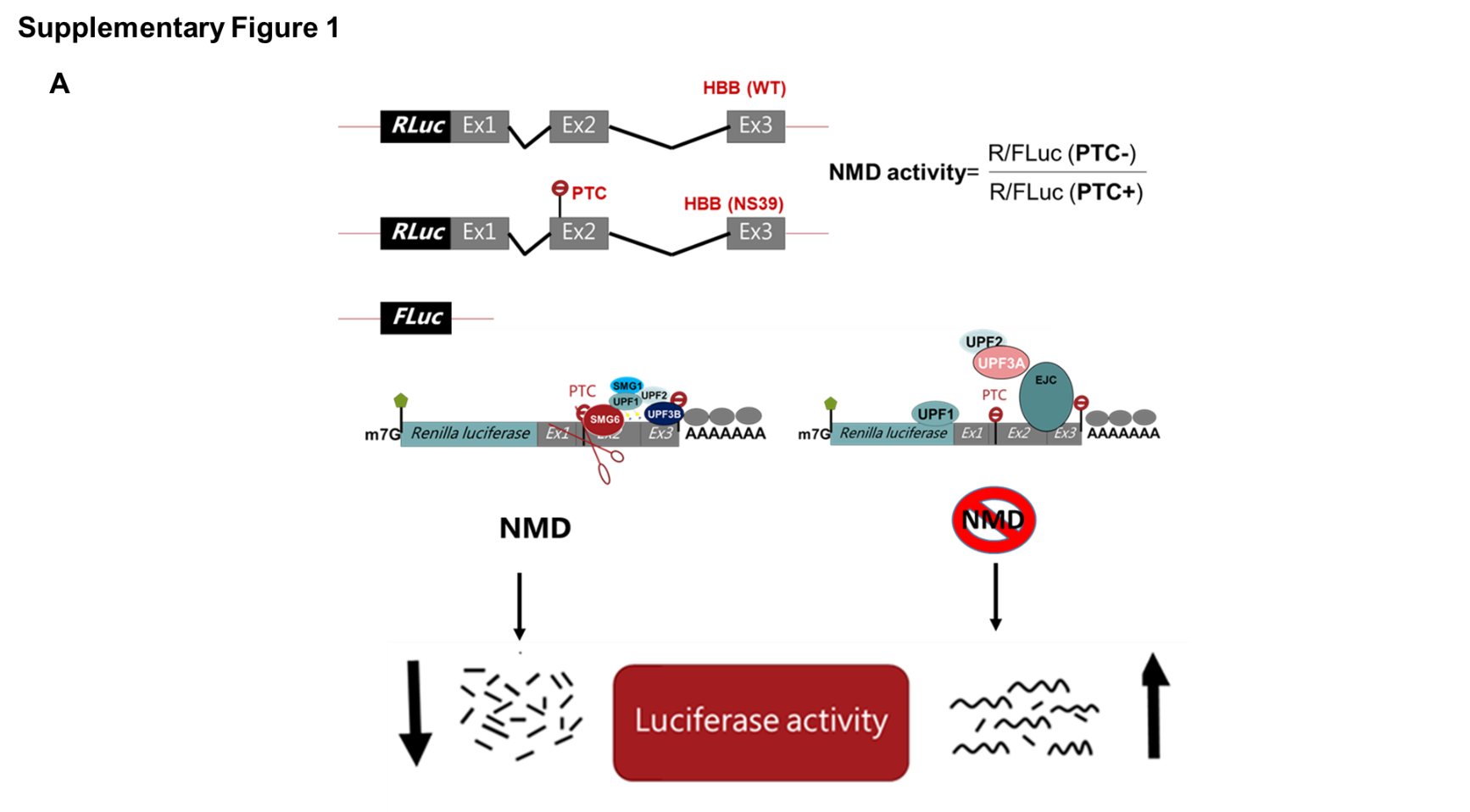

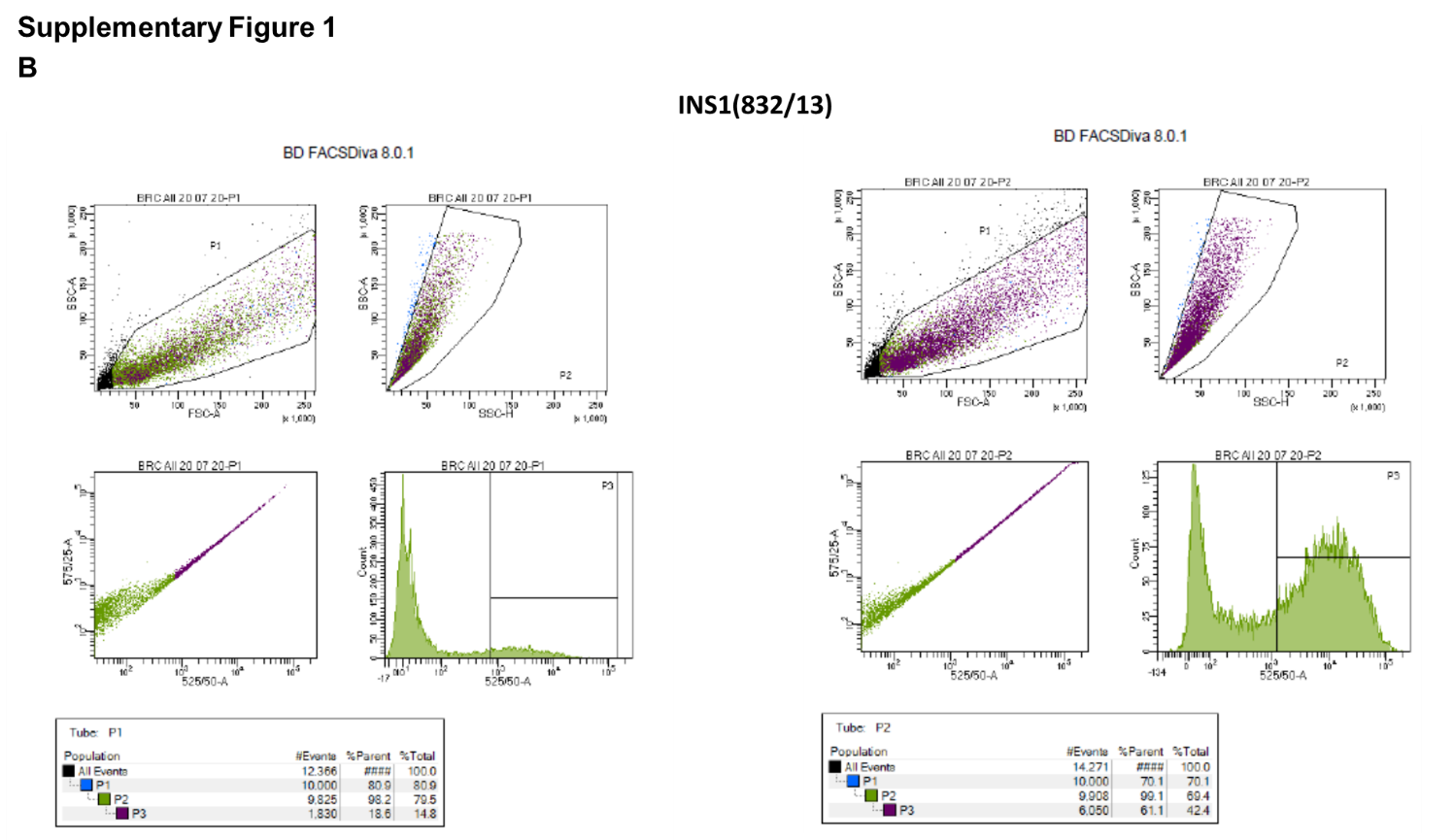

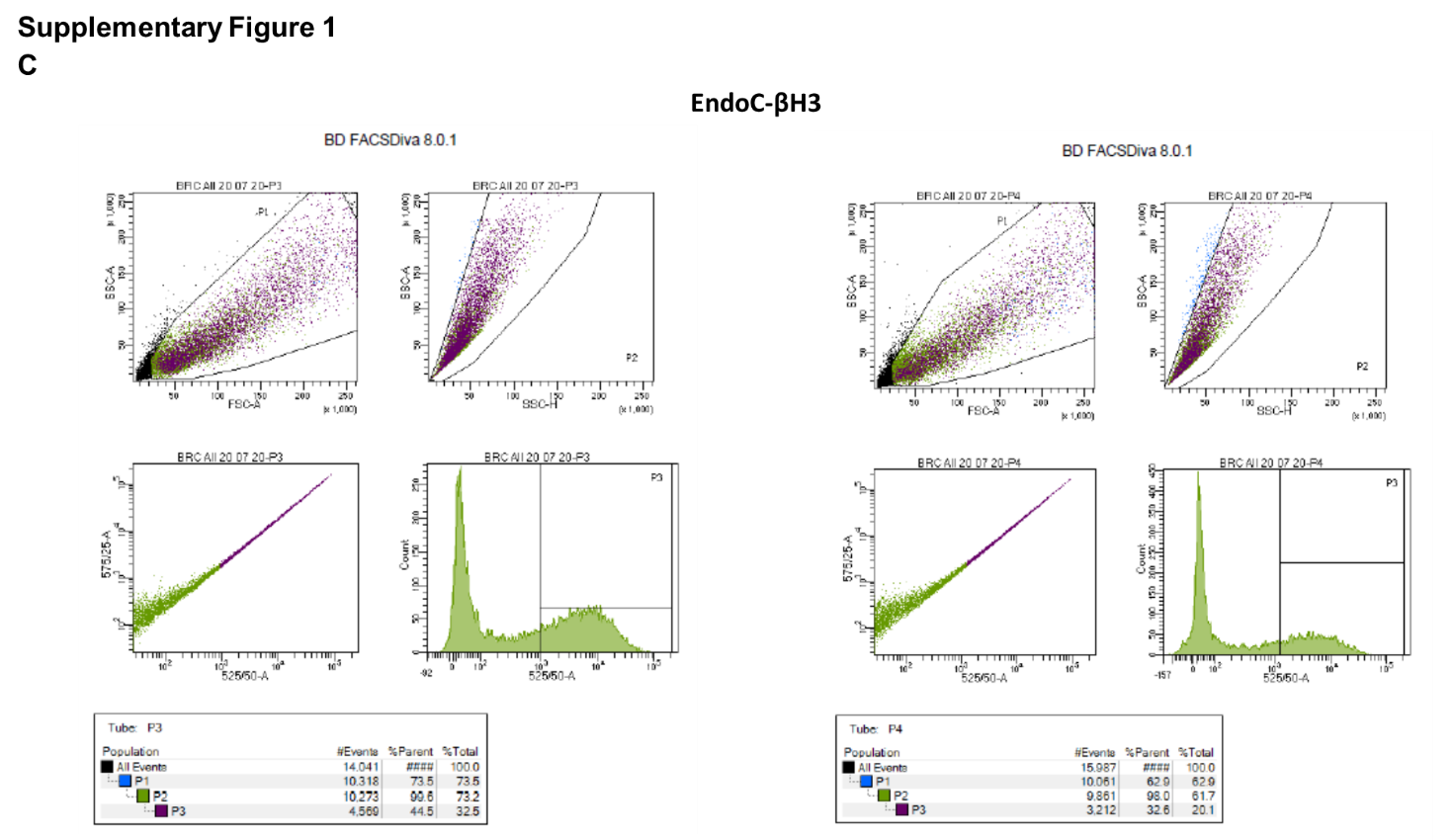

### Supplementary Figure 1. Measurement of NMD activity and transfection efficiencies in pancreatic β-cells

A. A schematic representation of luciferase-based NMD activity reporter which expresses either a wild-type (WT or namely PTC-) or a PTC-containing mutation (NS39 or namely PTC+) of *Haemoglobin-β (HBB)* gene, designated: HBB(PTC-) and HBB(PTC+), respectively, fused to the *Renilla* luciferase (*RLuc*) gene, and an empty plasmid expressing a *Firefly* luciferase (*FLuc*) as transfection control efficiency.

B-C. INS1(832/13) (B), EndoC-βH3 (C) cells were transfected by a GFP expressing plasmid and transfection efficiency was measured by FACS analysis 24 hrs post-transfection as explained in detail in the supplementary methods.

### Supplementary Figure 2

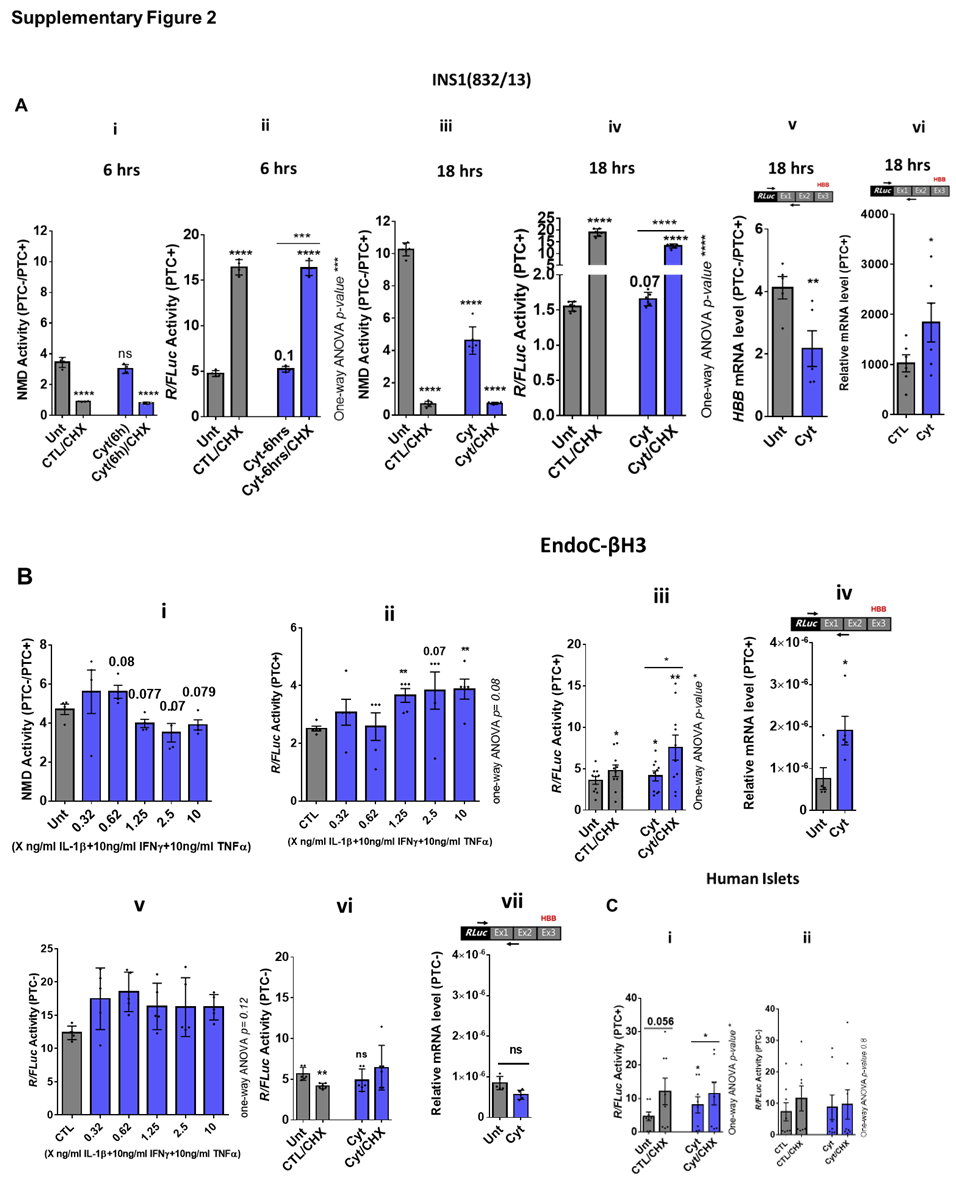

### Supplementary Figure 2. Inflammatory cytokines suppress NMD activity in β-cells

A-C. INS1(832/13) (A), EndoC-βH3 (B) cells and dispersed human islet cells (C) were co-transfected with HBB(PTC-) and or HBB(PTC+) and the *Firefly* plasmid and exposed to cytokines combination and or PBS as untreated (Unt) simultaneously with or without 10 µg/ml Cycloheximide (CHX) as an inhibitor of NMD activity.

A (i-iv). Luciferase activity (*R/FLuc* activity of PTC+) was measured in the lysate of the transfected INS1(832/13) cells exposed to cytokines combination (Cyt; 150 pg/mL IL-1β + 0.1 ng/mL IFNγ+0.1 ng/mL TNFα) for 6 hrs and 18 hrs as denoted on top.

B (i-vi). Luciferase activity (*R/FLuc* activity of PTC+) was measured in the lysate of the transfected EndoC-βH3 cells exposed to cytokines combination (Cyt; X ng/mL IL-1β + 10 ng/mL IFNγ+10 ng/mL TNFα) (right) and (Cyt; 3 ng/mL IL-1β + 10 ng/mL IFNγ+10 ng/mL TNFα) (left) for 18 hrs.

A (v and vi) and B (iv). mRNA level of Renilla-HBB(PTC+) fused gene and *Firefly* gene in the transfected INS1(832/13) (A-v and vii) and EndoC-βH3 (B-iv) cells was quantified by RT-qPCR using specific primers extending the junction of exons 1 and 2 of the *HBB* gene, and *Renilla* gene, or only *Firefly* gene and normalised to actin and tubulin, respectively.

C (i-ii). Luciferase activity (*R/FLuc* activity of PTC+) was measured in the lysate of the transfected dispersed human islet cells exposed to cytokines combination (Cyt; 3 ng/mL IL-1β + 10 ng/mL IFN-γ+10 ng/mL TNFα) (left) for 18 hrs.

The data are means ± SEM of N=6. The symbol * indicates the Bonferroni-corrected paired t-test values of treatments versus untreated (Unt), cytokines (Cyt) that is, otherwise, designated by a line on top of the bars (A-C). * ≤ 0.05, ** ≤ 0.01, *** ≤0.001, **** ≤ 0.0001. ns: non-significant.

### Supplementary Figure 3

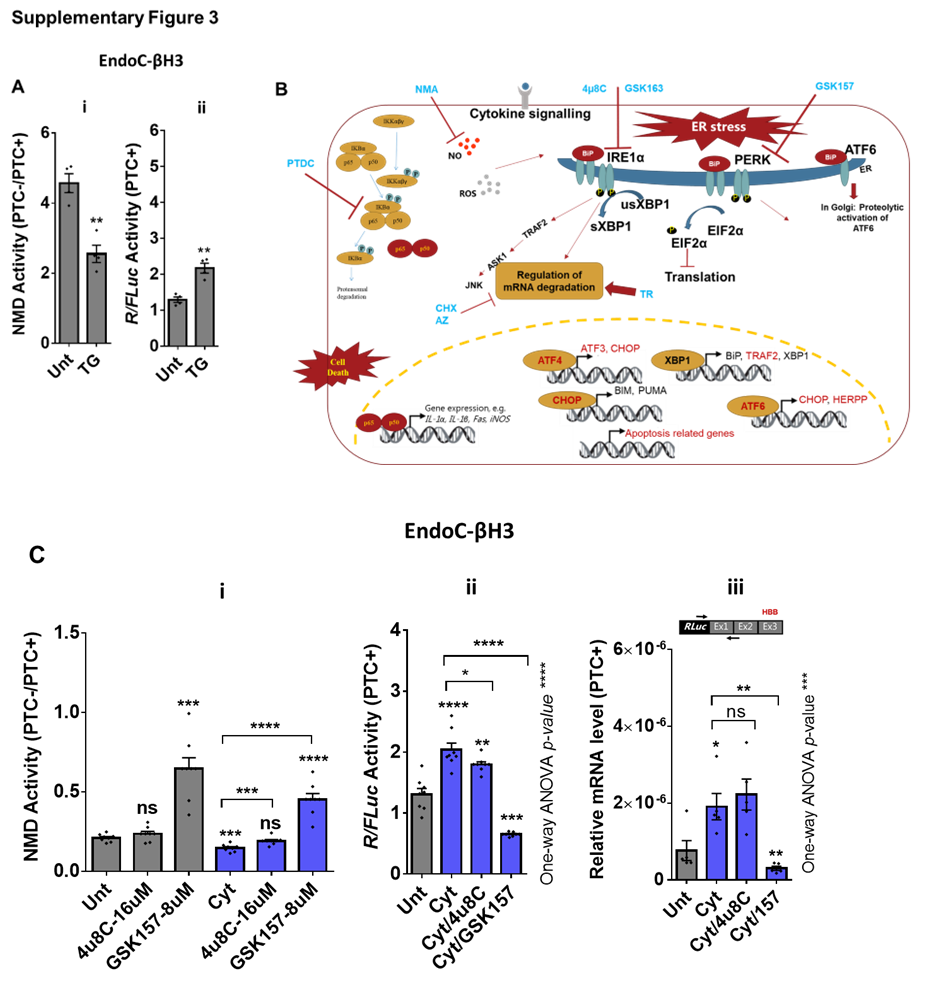

### Supplementary Figure 3. ER stress is involved in the inflammatory cytokines-induced suppression of NMD activity in β-cells

A. EndoC-βH3 cells were co-transfected with HBB(PTC-) and or HBB(PTC+) and *Firefly* plasmids and exposed to PBS as untreated (Unt) or 1µM Thapsigargin (TG) or for 18 hrs. Luciferase activity (*R/FLuc* activity of PTC+) was measured in the lysate of EndoC-βH3 cells transfected with HBB(PTC+) exposed to PBS as untreated (Unt) or 1µM Thapsigargin (TG), an ER stress inducer as explained in the methods.

B. a scheme of using chemical inhibition of riboendonuclease and kinase activities of IRE1α with 4µ8C and GSK2850163 (GSK163), respectively, PERK activation with GSK2656157 (GSK157), NF-κB with Ammonium pyrrolidinedithiocarbamate (PDTC) and iNOS with NG-methyl-L-arginine (NMA), and NMD activity with 10 µg/ml Cycloheximide (CHX) and 5-Azacytidine (AZ), and of NMD activation with Tranilast (TR). Data for CHX, AZ and TR are not shown here.

C. EndoC-βH3 cells were co-transfected with HBB(PTC-) and or HBB(PTC+) and *Firefly* plasmids and exposed to PBS as untreated (Unt), cytokines combination (Cyt; 3 ng/mL IL-1β + 10 ng/mL IFNγ+10 ng/mL TNFα) alone, and or simultaneously with 16 µM of 4µ8C, an endoribonuclease inhibitor of IRE1α, and or 8 µM of GSK2656157 (GSK157) for 18 hrs. C(i and ii). Luciferase activity (*R/FLuc* activity of *PTC*+) was measured in the lysate of the EndoC-βH3 cells as explained in the methods. C(ii). mRNA level of *Renilla-*HBB(PTC+) fused gene and *Firefly* gene in the EndoC-βH3 cells was quantified by RT-qPCR using specific primers extending the junction of exons 1 and 2 of the *HBB* gene, and *Renilla* gene, or only *Firefly* gene and normalised to tubulin.

The data are means ± SEM of N=6. The symbol * indicates the Bonferroni-corrected paired t-test values of treatments versus untreated (Unt) (A and C) or cytokines (Cyt) that is, otherwise, designated by a line on top of the bars (C). * ≤ 0.05, ** ≤ 0.01, *** ≤0.001, **** ≤ 0.0001.

### Supplementary Figure 4

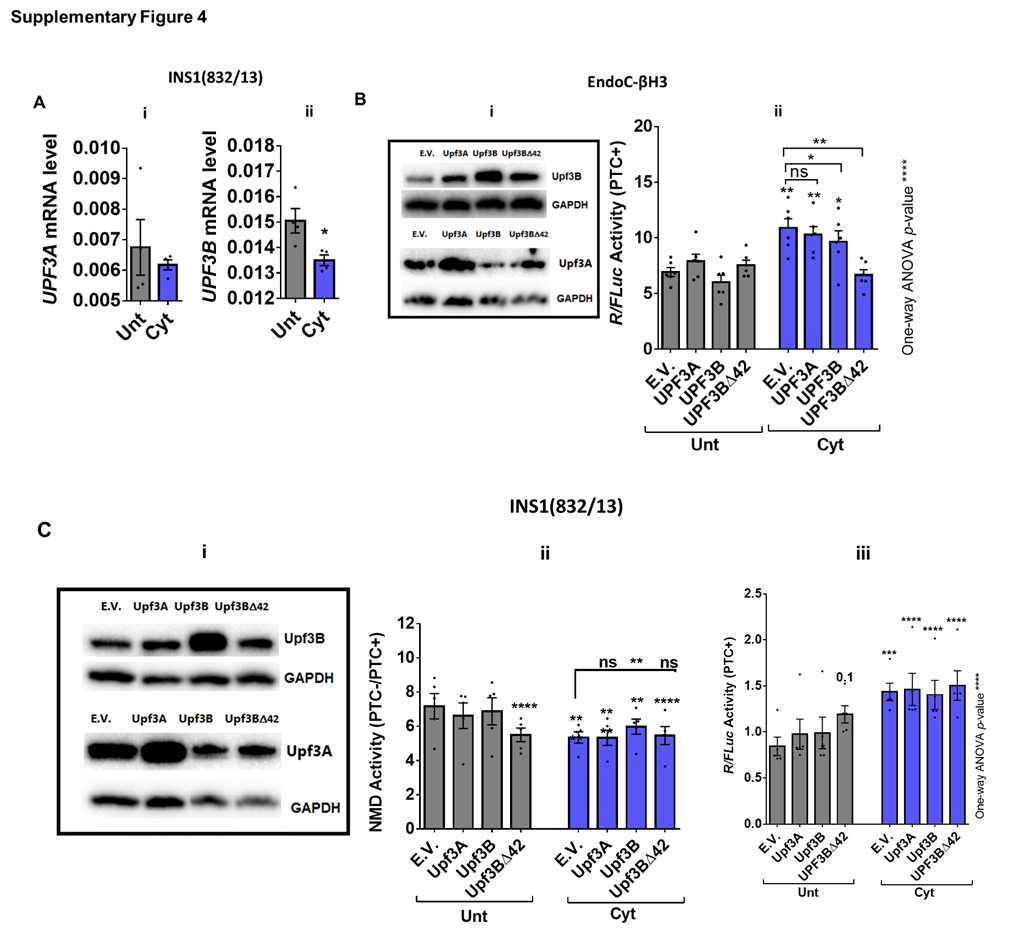

### Supplementary figure 4. Inflammatory cytokines-induced suppression of NMD activity is also driven by UPF3B downregulation in β-cells.

INS1(832/13) and EndoC-βH3 cells were co-transfected with empty vector (E.V.), UPF3A, UPF3B and or UPF3BΔ42 (dominant negative of UPF3B) plasmids, and with HBB(PTC-) and or HBB(PTC+), and *Firefly* plasmids and exposed to cytokine combination (Cyt; 150 pg/mL IL-1β + 0.1 ng/mL IFNγ+ 0.1 ng/mL TNFα) and (Cyt; 3 ng/mL IL-1β + 10 ng/mL IFN-γ+10 ng/mL TNFα), respectively, for 18 hrs.

A (i and ii). mRNA level of *Upf3A* and *Upf3B* genes in INS1(832/13) cells was quantified by RT-qPCR and normalised to actin mRNA.

B (ii). Luciferase activity (*R/FLuc* activity of PTC+) was measured in the lysate of the transfected EndoC-βH3 cells as explained in the methods. The overexpression of Upf3A and Upf3B proteins was examined in the cell lysate (B-i) by Western blot analysis.

C (ii). Luciferase activity was measured in the lysate of INS1 (812/13) cells transfected with HBB (PTC-) and or HBB(PTC+) represented as NMD activity calculated by dividing luciferase activity of HBB (PTC-)/HBB (PTC+) as explained in the methods. The overexpression of Upf3A and Upf3B proteins was examined in the INS1 (832/13) lysate (C-i) by Western blot analysis.

The data are means ± SEM of N=6. The symbol * indicates the Bonferroni-corrected paired t-test values of treatments versus untreated E.V. (Unt) (A-C) or cytokines (Cyt)-treated E.V. that is, otherwise, designated by a line on top of the bars (B-C). * ≤ 0.05, ** ≤ 0.01, *** ≤0.001, **** ≤ 0.0001. ns: non-significant.

### Supplementary Figure 5

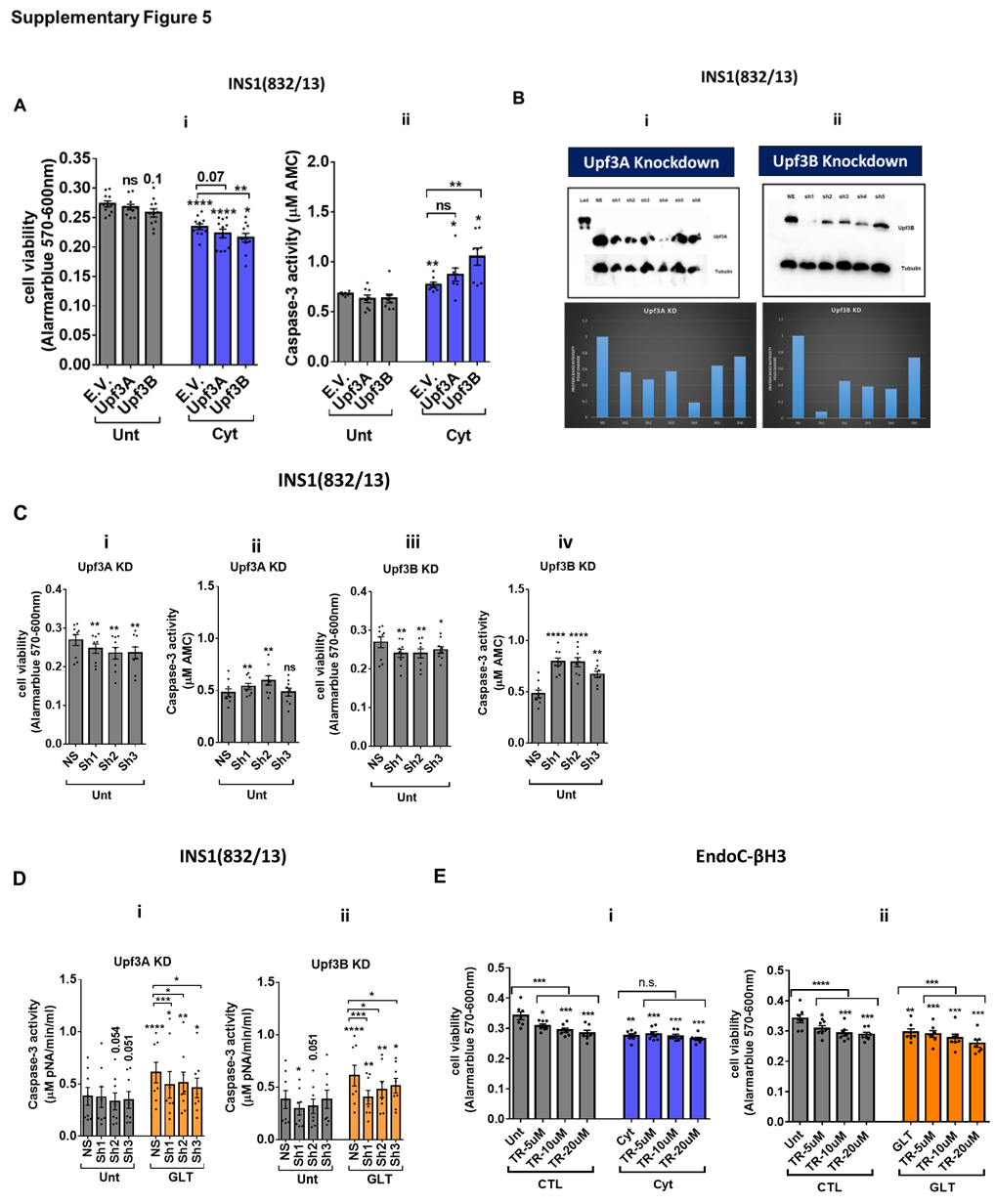

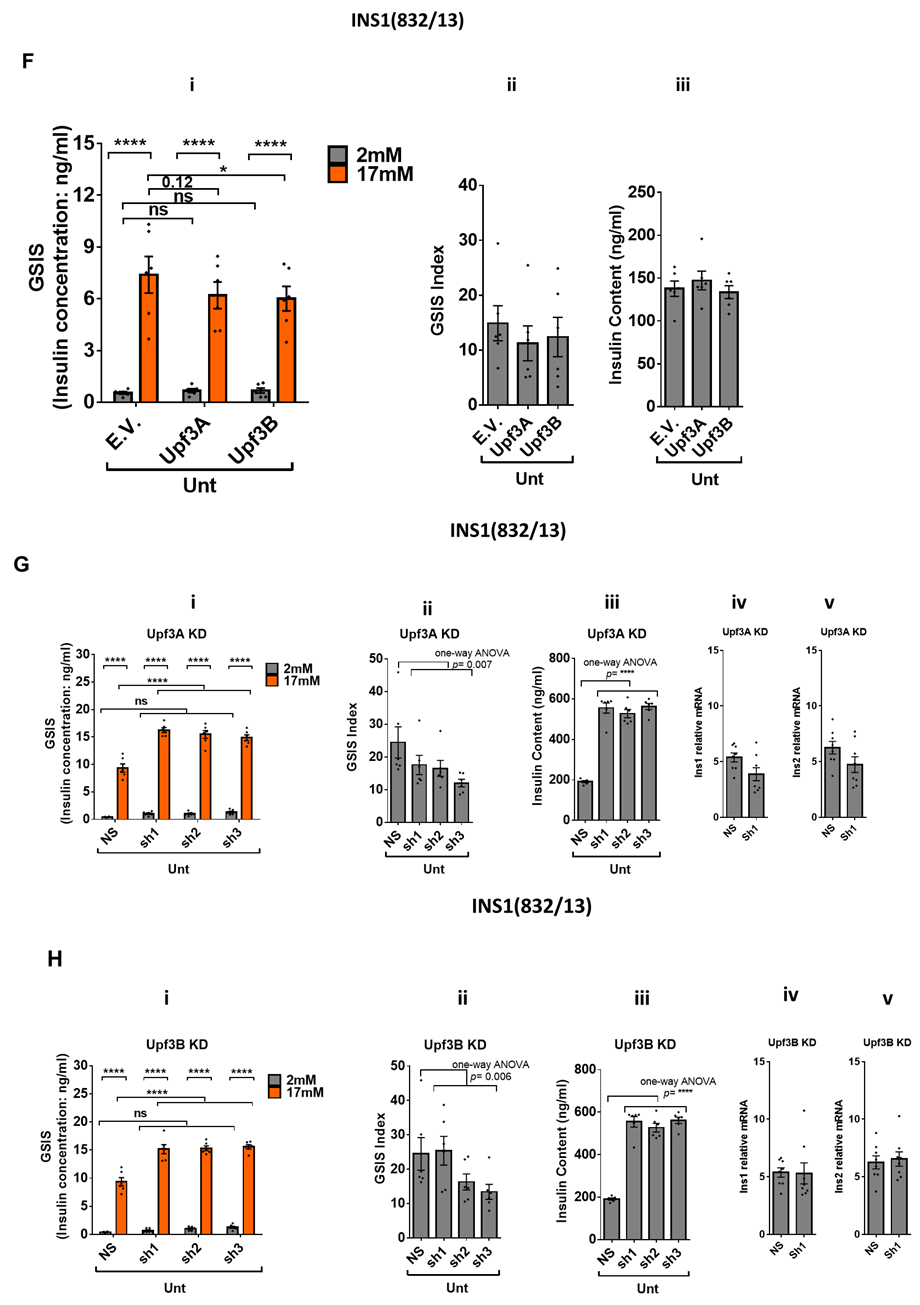

### Supplementary figure 5. Upf3A/B knockdown deteriorates basal cell viability associated with increased insulin content, whereas reduces glucolipotoxicity-mediated cell death in INS1 (832/13) cells.

A (i and ii). INS1(832/13) cells were co-transfected with empty vector (E.V.), Upf3A and or Upf3B plasmids and then, with HBB(PTC-) and or HBB(PTC+) and exposed to cytokines combination (Cyt; 150 pg/mL IL-1β + 0.1 ng/mL IFNγ+ 0.1 ng/mL TNFα) for 24 hrs. Cell viability was measured by Alamarblue (i) and caspase-3 activity (µM AMC) (ii) assays (N=6).

B-D. INS1(832/13) cell lines with the most efficient stable knock-down of UPF3A and UPF3B (three shRNAs) and non-silencing shRNA control (NS) qualified by quantitative WB (B-i and ii) were treated with PBS (C-D), and or glucolipotoxicity (D). The protein band intensity of the WB was quantified using ImageJ and represented as graph (B-bottom). Cell viability was measured by Alamarblue (C-i and iii) and caspase-3 activity (µM AMC) (C-ii and iv) assays in untreated cells, but only caspase-3 activity for glucolipotoxicity effect (D-i and ii) (N=6). The caspase-3 activity (µM pNA/min/ml) for glucolipotoxicity effect was measured using colorimetric caspase-3 activity assay kit according to manufacturer’s protocol.

E. EndoC-βH3 cells were exposed to cytokines combination (Cyt; 3 ng/mL IL-1β + 10 ng/mL IFNγ+10 ng/mL TNFα) (E-i) and or glucolipotoxicity (E-ii) alone or with Tranilast at given concentrations for three days. Cell viability was measured by Alamarblue assay (N=6).

F. INS1(832/13) cells were co-transfected with empty vector (E.V.), UPF3A and or UPF3B plasmids and experimented for glucose-stimulated insulin secretion (GSIS) (i) and insulin contents (iii). Insulin concentration (ng/ml) was measured by insulin ultra-sensitive assay (N=6). GSIS index (ii) was calculated by dividing insulin concentration measured in the treatments of 17mM by 2mM glucose.

G-H. INS1(832/13) cell lines with the most efficient stable knock-down of UPF3A (G) and UPF3B (H) (three shRNAs) and non-silencing shRNA control (NS) experimented for glucose-stimulated insulin secretion (GSIS) (Subpanels i) and insulin contents (Subpanels iii). Insulin concentration (ng/ml) was measured by insulin ultra-sensitive assay (N=6). GSIS index (Subpanels ii) was calculated by dividing insulin concentration measured in the treatments of 17 mM by 2 mM glucose. The levels of insulin 1 (Ins1) and insulin 2 (Ins2) mRNAs were measured by RT-qPCR and normalised to HPRT1 in the UPF3A (G-iv and v) and UPF3B KD (H-iv and v) cells.

The symbol * indicates the Bonferroni-corrected paired t-test values of treatments versus untreated E.V. (Unt), untreated nonsense control (NS), or cytokines (Cyt)- or GLT-treated conditions that is, otherwise, designated by a line on top of the bars, or corresponding low/high glucose untreated E.V. or NS controls which is designated by lines on the top of the bars (B). * ≤ 0.05, ** ≤ 0.01, *** ≤0.001, **** ≤ 0.0001. ns: non-significant.

### Supplementary Figure 6

### Supplementary figure 6. UPF2 knockdown boosts cytokines suppression of NMD activity in EndoC-βH3 cells

EndoC-βH3 cell lines with the most efficient stable knock-down of UPF2 (three shRNAs: sh1, sh4 and sh5) and a non-silencing shRNA control (NS) qualified by quantitative WB (A) were co-transfected with HBB(PTC-) and or HBB(PTC+) and *Firefly* plasmids and exposed to PBS as untreated (Unt), cytokines combination (Cyt; 3 ng/mL IL-1β + 10 ng/mL IFNγ+10 ng/mL TNFα). The protein band intensity (A-i) of the WB was quantified using ImageJ as represented as graph (A-ii). Luciferase activity (B) was measured in the lysate of the transfected cells represented as NMD activity (B-i) calculated by dividing luciferase activity of HBB(PTC-)/HBB(PTC+) and luciferase activity of HBB(PTC+) (*R/FLuc* activity of *PTC*+; B-ii) as explained in the methods (N=5).

The symbol * indicates the Bonferroni-corrected paired t-test values of treatments versus untreated (Unt) NS control or cytokines, or otherwise, designated by lines on the top of the bars (B). * ≤ 0.05, ** ≤ 0.01, *** ≤0.001, **** ≤ 0.0001.

### Supplementary Figure 7

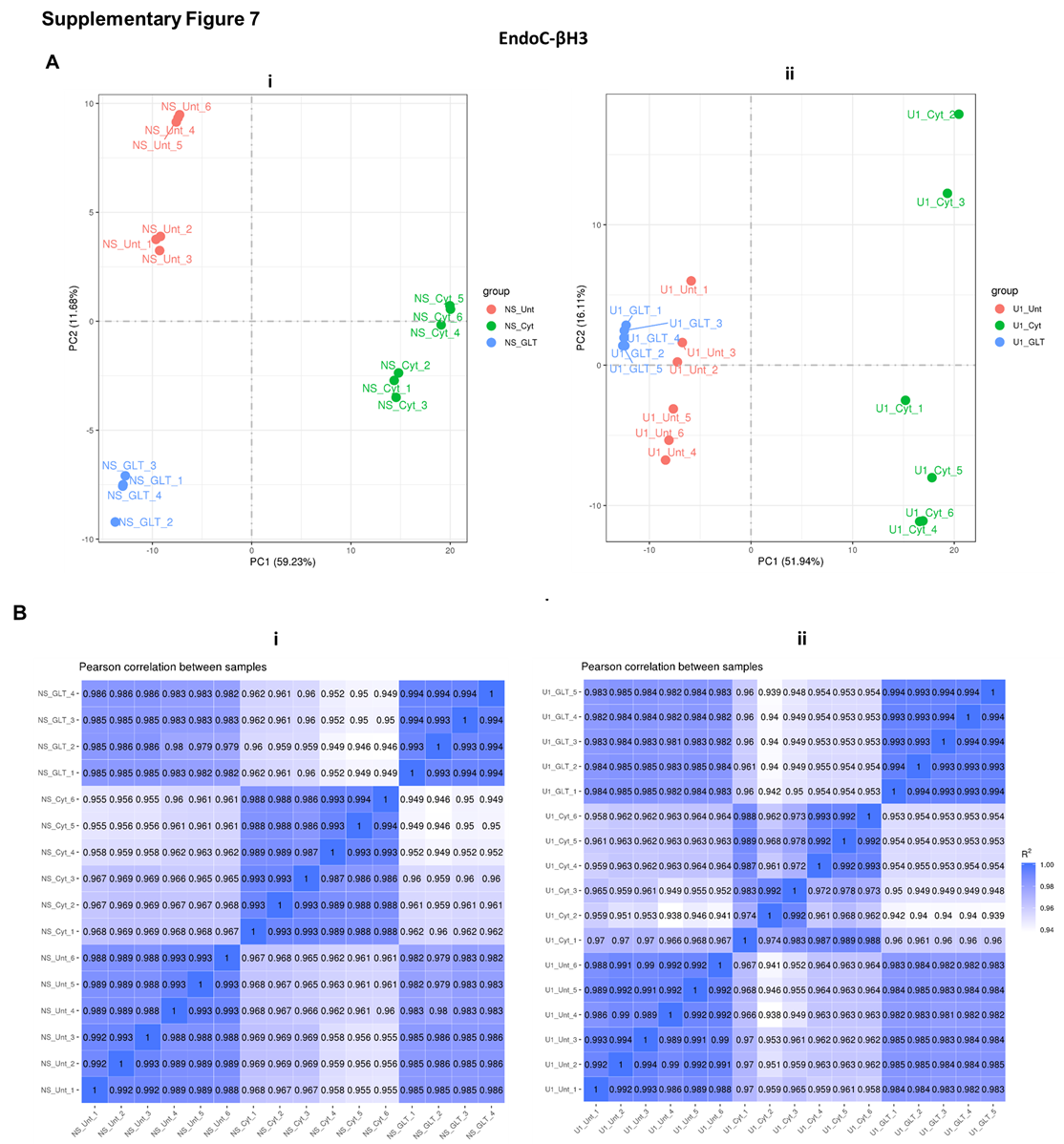

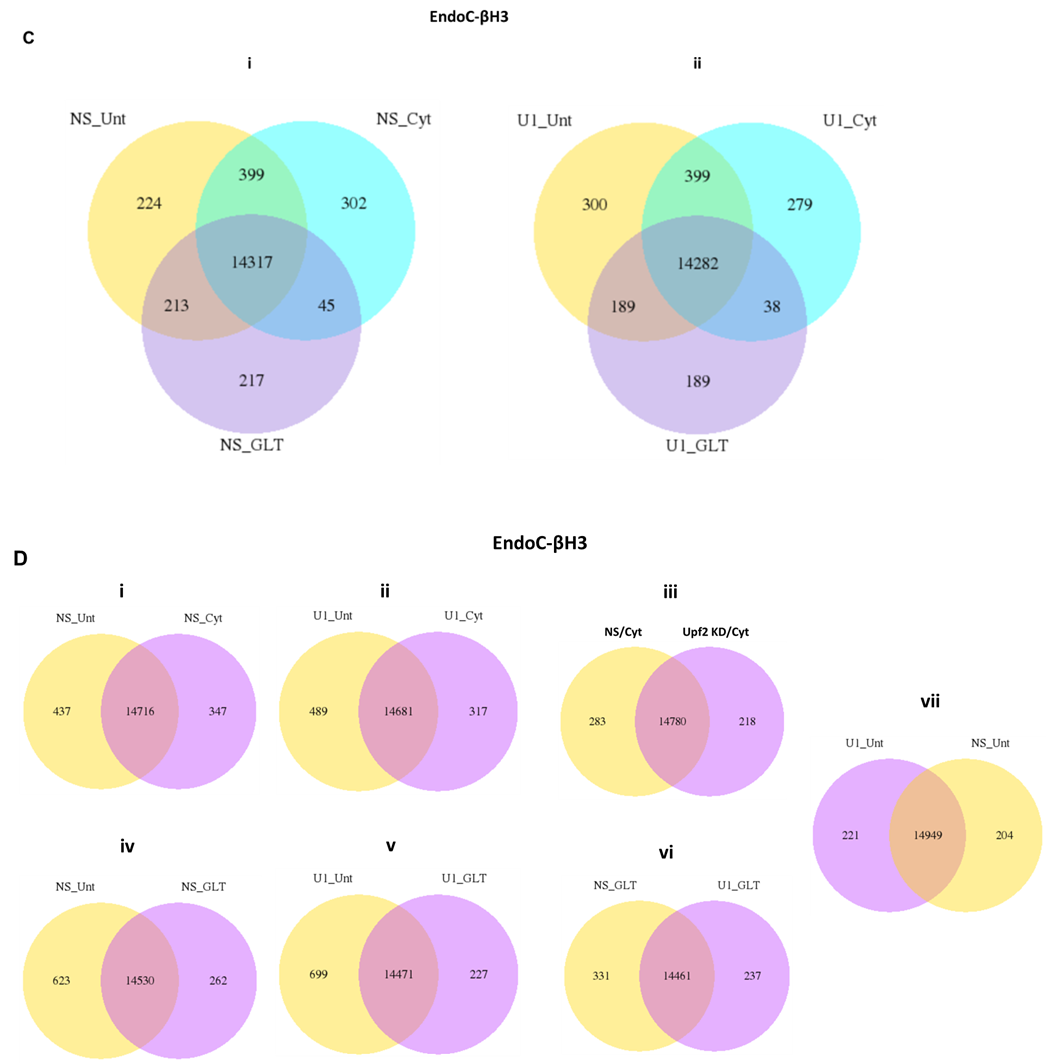

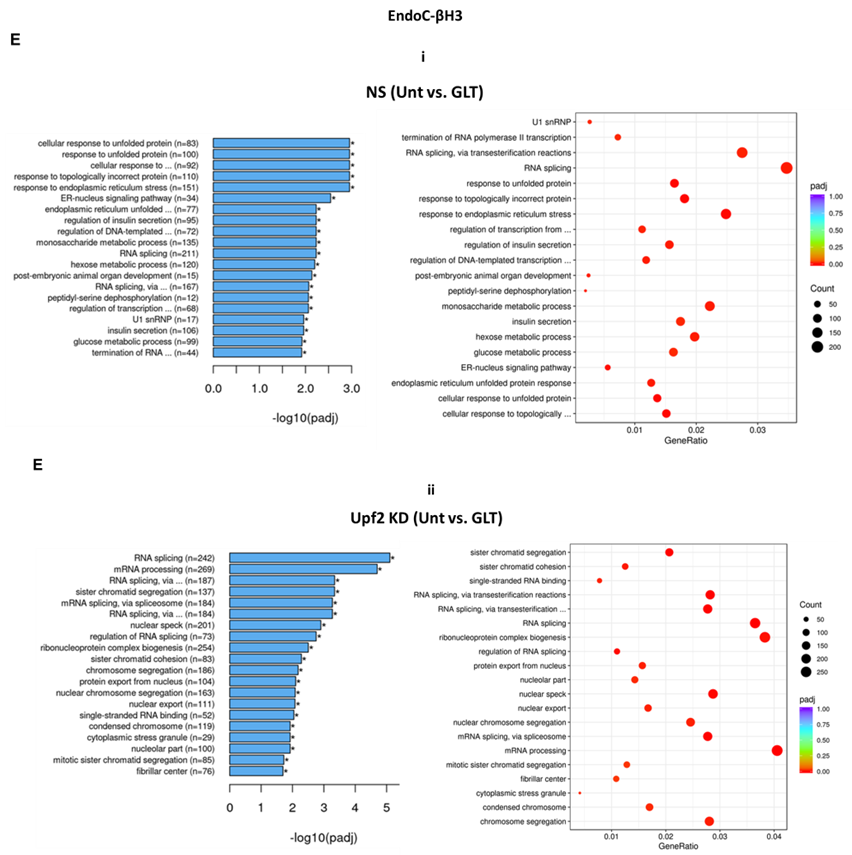

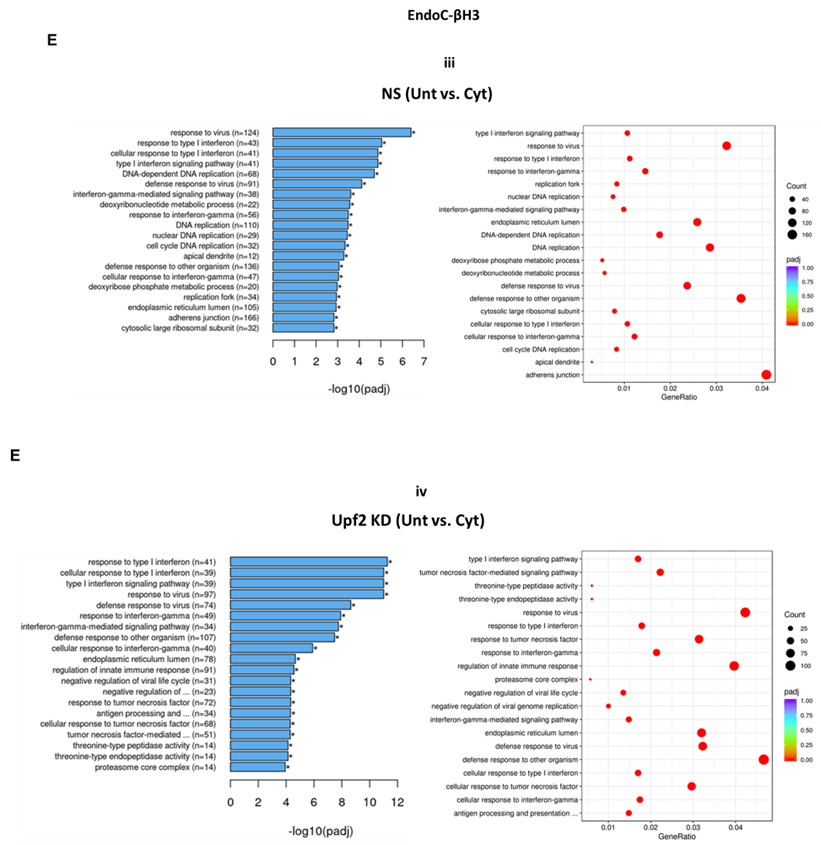

### Supplementary figure 7. UPF2 knockdown differentially implicates in cytokines and glucolipotoxicity deregulation of EndoC-βH3 transcripts.

EndoC-βH3 cell lines with the most efficient stable knock-down of UPF2 (U1; shRNA-1) and non-silencing control (NS) were exposed to PBS as untreated (Unt), cytokines combination (Cyt; 3 ng/mL IL-1β + 10 ng/mL IFNγ+10 ng/mL TNFα) and or glucolipotoxicity (GLT; 0.5 mM Palmitate+25 mM glucose). Total RNA was extracted from the treated cells, cDNA library was made and sequenced using Hiseq platform under contracted company’s manual. RNA-seq datasets including NS/CTL (N=6), NS/Cyt (N=6), NS/GLT (N=4), U1/CTL (N=6), U1/Cyt (N=6) and U1/GLT (N=5) were analysed through the pipeline described in the methods.

A. Principal component analysis (PCA) of transcripts regulated under Unt, Cyt and or GLT conditions in the samples from NS (i) and UPF2 KD (U1) (ii) cell lines.

B. Pearson correlation between the samples from Unt, Cyt and GLT exposed NS (i) and UPF2 KD (U1) (ii) cell lines.

C-D. Venn diagram presentation of the number of commonly and differentially regulated transcripts by Cyt or GLT as indicated in the figures.

E. Top GEA-identified transcripts regulated by GLT (i and ii) Cyt (iii and iv) compared to untreated in UPF2 KD vs. NS control cells. The expression level is shown as log2 (adjusted *p*-value <0.05).

### Supplementary Figure 8

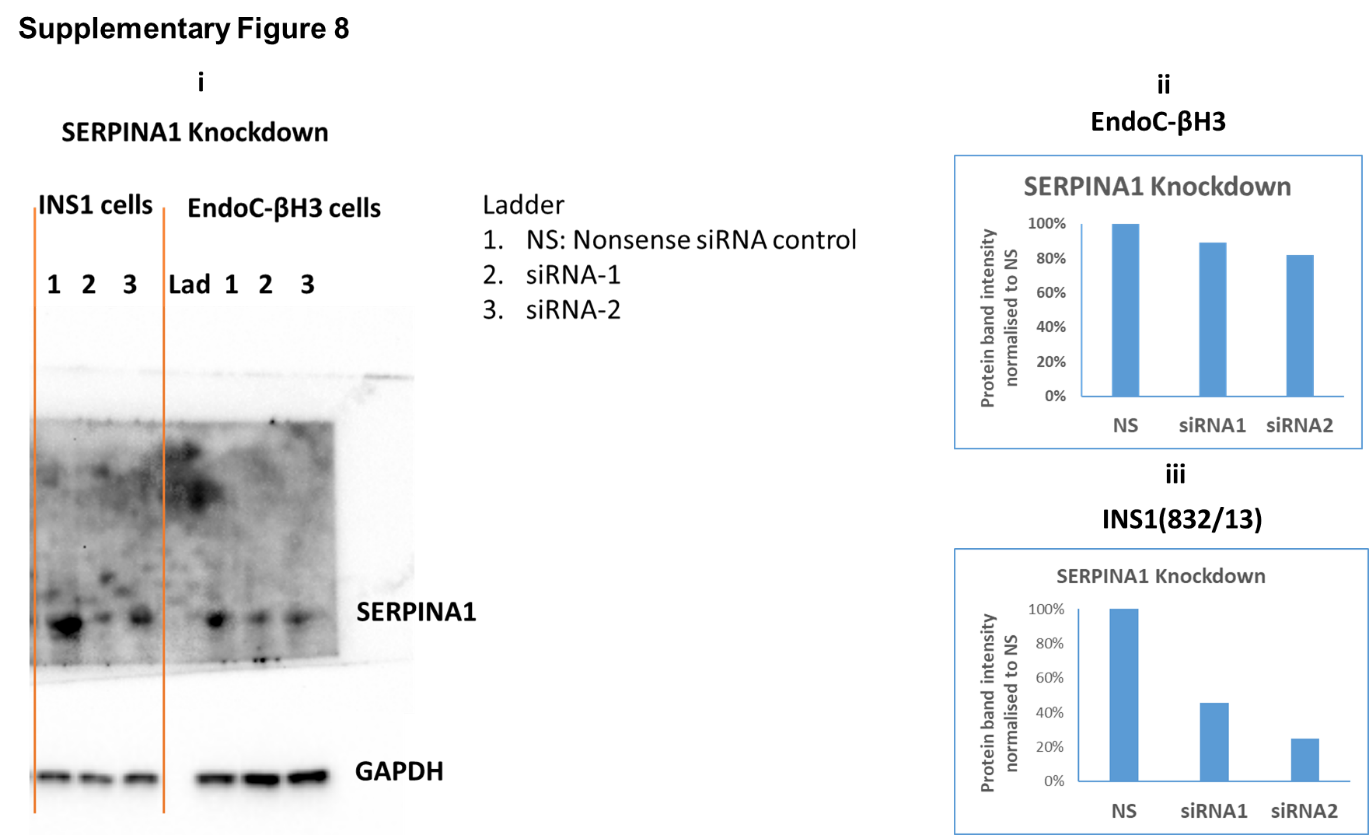

### Supplementary Figure 8. SERPINA1 knockdown deteriorates cytokine cytotoxicity for viability and glucose-stimulated insulin secretion index in INS1(832/13) cells.

EndoC-βH3 and INS1(832/13) cells were transfected with siRNAs against SERPINA1 (two species-specific siRNAs for each cell type) and a non-silencing siRNA control (NS), incubated for 24 hours. The knockdown efficiency was checked using quantitative WB (i) and the protein band intensity of the WB was quantified using ImageJ as represented as graph (ii and iii) as described in the methods.

### Supplementary Figure 9

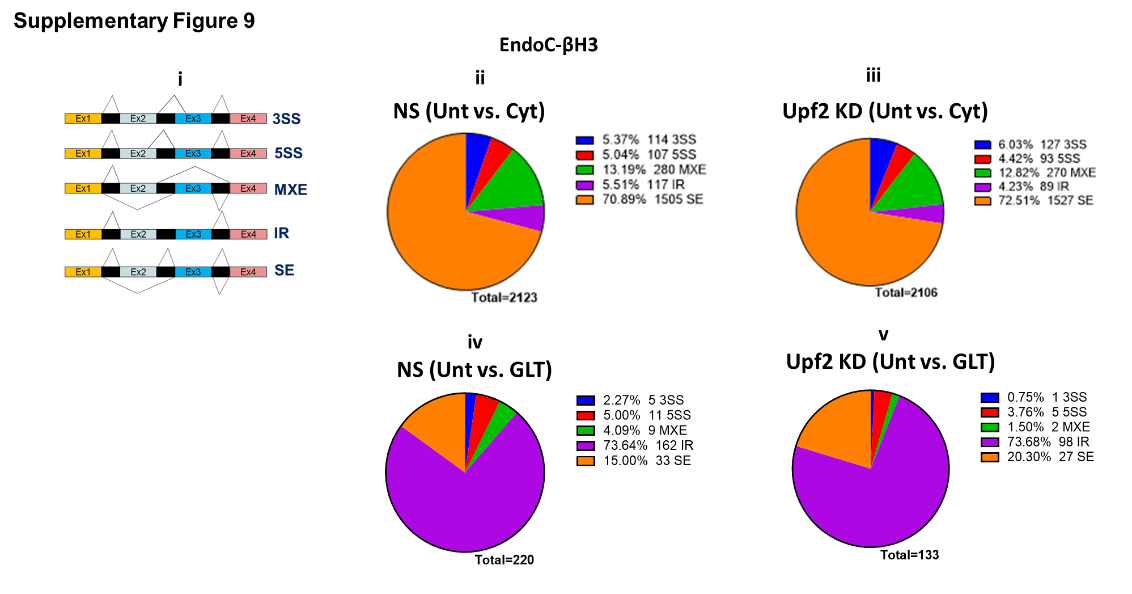

### Supplementary Figure 9. UPF2 knockdown potentiates the skipped exon rate of alternatively spliced transcript isoforms increased by cytokines, but not glucolipotoxicity, in EndoC-βH3.

EndoC-βH3 cell lines with the most efficient stable knock-down of UPF2 (U1; shRNA-1) and non-silencing control (NS) were exposed to PBS as control or untreated (Unt), cytokines combination (Cyt; 3 ng/mL IL-1β + 10 ng/mL IFNγ+10 ng/mL TNFα) and or glucolipotoxicity (GLT; 0.5 mM Palmitate+25 mM glucose). Total RNA was extracted from the treated cells, cDNA library was made and sequenced using Hiseq platform under contracted company’s manual. RNA-seq datasets including NS/CTL (N=6), NS/Cyt (N=6), NS/GLT (N=4), U1/CTL (N=6), U1/Cyt (N=6) and U1/GLT (N=5) were analysed through the pipeline performed by the company (Novogene, Cambridge, England). Alternative splicing events including alternative 3’ splice site (3SS), alternative 5’ splice site (3SS), mutually exclusive exons (MXE), intron retention (IR) and skipped exon (SE) driven by cytokines or GLT versus untreated in the NS control and Upf2 KD EndoC-βH3 cells were analysed using the software rMATS as described in the methods.

### References

1.	Benazra M, Lecomte MJ, Colace C, Muller A, Machado C, Pechberty S, et al. A human beta cell line with drug inducible excision of immortalizing transgenes. Mol Metab. 2015;4(12):916-25.

2.	Doliba NM, Liu Q, Li C, Chen J, Chen P, Liu C, et al. Accumulation of 3-hydroxytetradecenoic acid: Cause or corollary of glucolipotoxic impairment of pancreatic beta-cell bioenergetics? Mol Metab. 2015;4(12):926-39.

3.	Prause M, Christensen DP, Billestrup N, Mandrup-Poulsen T. JNK1 protects against glucolipotoxicity-mediated beta-cell apoptosis. PLoS One. 2014;9(1):e87067.

4.	Sekine N, Fasolato C, Pralong WF, Theler JM, Wollheim CB. Glucose-induced insulin secretion in INS-1 cells depends on factors present in fetal calf serum and rat islet-conditioned medium. Diabetes. 1997;46(9):1424-33.

5.	Bugliani M, Syed F, Masini M, Marselli L, Suleiman M, Novelli M, et al. Direct effects of rosuvastatin on pancreatic human beta cells. Acta Diabetol. 2013;50(6):983-5.

6.	Boelz S, Neu-Yilik G, Gehring NH, Hentze MW, Kulozik AE. A chemiluminescence-based reporter system to monitor nonsense-mediated mRNA decay. Biochem Biophys Res Commun. 2006;349(1):186-91.

7.	Neu-Yilik G, Raimondeau E, Eliseev B, Yeramala L, Amthor B, Deniaud A, et al. Dual function of UPF3B in early and late translation termination. EMBO J. 2017;36(20):2968-86.

8.	Lever J, Krzywinski M, Altman N. Principal component analysis. Nature Methods. 2017;14(7):641-2.

9.	Andersen CL, Jensen JL, Orntoft TF. Normalization of real-time quantitative reverse transcription-PCR data: a model-based variance estimation approach to identify genes suited for normalization, applied to bladder and colon cancer data sets. Cancer Res. 2004;64(15):5245-50.

10.	Ghiasi SM, Krogh N, Tyrberg B, Mandrup-Poulsen T. The No-Go and Nonsense-Mediated RNA Decay Pathways Are Regulated by Inflammatory Cytokines in Insulin-Producing Cells and Human Islets and Determine beta-Cell Insulin Biosynthesis and Survival. Diabetes. 2018;67(10):2019-37.
